# Interpretable PROTAC degradation prediction with structure-informed deep ternary attention framework

**DOI:** 10.1101/2024.11.05.622005

**Authors:** Zhenglu Chen, Chunbin Gu, Shuoyan Tan, Xiaorui Wang, Yuquan Li, Mutian He, Ruiqiang Lu, Shijia Sun, Chang-Yu Hsieh, Xiaojun Yao, Huanxiang Liu, Pheng-Ann Heng

**Affiliations:** Lanzhou University; Macao Polytechnic University; The Chinese University of Hong Kong; Zhejiang University

**Author notes:** These authors contributed equally to this work (Zhenglu Chen, Chunbin Gu).

## Abstract

Proteolysis Targeting Chimeras (PROTACs) are heterobifunctional ligands that form ternary complexes with Protein Of Interests (POIs) and E3 ligases, exploiting the ubiquitin-proteasome system to degrade disease-associated proteins, promising to drug the ‘undruggable’. While PROTAC research primarily relies on costly and time-consuming wet experimental approaches, deep learning offers a promising avenue to accelerate development and reduce expenses. However, existing deep learning methods for PROTAC degradation prediction often overlook the significance of hierarchical molecular representation and protein structural information, hindering effective data modeling. Moreover, their black-box nature limits the interpretability of computational outcomes, failing to provide intuitive insights into substructure interactions within the PROTAC system. This study introduces *PROTAC-STAN*, a structure-informed deep ternary attention network (STAN) framework for interpretable PROTAC degradation prediction. *PROTAC-STAN* represents PROTAC molecules across atom, molecule, and property hierarchies and incorporates structure information for POIs and E3 ligases using a protein language model infused with structural data. Furthermore, it simulates interactions among three entities at the atom and amino acid levels via a novel ternary attention network tailored for the PROTAC system, providing unprecedented insights into the degradation mechanism. By integrating hierarchical PROTAC molecule representation, structural embedding of POI and E3 ligase, and ternary attention network modeling interactions, our approach substantially improves prediction accuracy by 10.95% while enabling significant model interpretability via atomic and residue level visualization of molecule and complex. Experiments on the refined public PROTAC dataset demonstrate that *PROTAC-STAN* outperforms state-of-the-art baselines in overall performance. The excellent performance of *PROTAC-STAN* is anticipated to establish it as a foundational tool in future research on PROTAC-related drugs, thereby accelerating the development of this field.

## 1 Introduction

Targeted Protein Degradation (TPD) has presented a promising approach to modern drug development in recent years [1–3]. Unlike traditional drug design methods that typically modulate protein function through inhibition or activation, TPD targets proteins for degradation by marking them as substrates for cellular degradation machinery, thus achieving therapeutic outcomes [1]. TPD holds the potential to address proteins considered ‘undruggable’ by traditional methods and may mitigate resistance caused by target pocket mutations [2, 4]. Moreover, it offers the advantage of reducing drug dosage and frequency, thereby minimizing adverse effects on patients [5]. Upon the foundation of TPD, Proteolysis Targeting Chimera (PROTAC) technology stands out as a compelling and novel therapeutic modality [3, 5]. As heterobifunctional ligands, PROTACs facilitate the interaction between POIs and E3 ligases, forming ternary complexes that exploit the ubiquitin-proteasome system (UPS) for selective degradation of disease-associated proteins [6] as illustrated in Figure 1. This strategy offers a more comprehensive approach compared to traditional small-molecule inhibitors or receptor antagonists, effectively eliminating pathological protein function [3].

**Fig 1.**
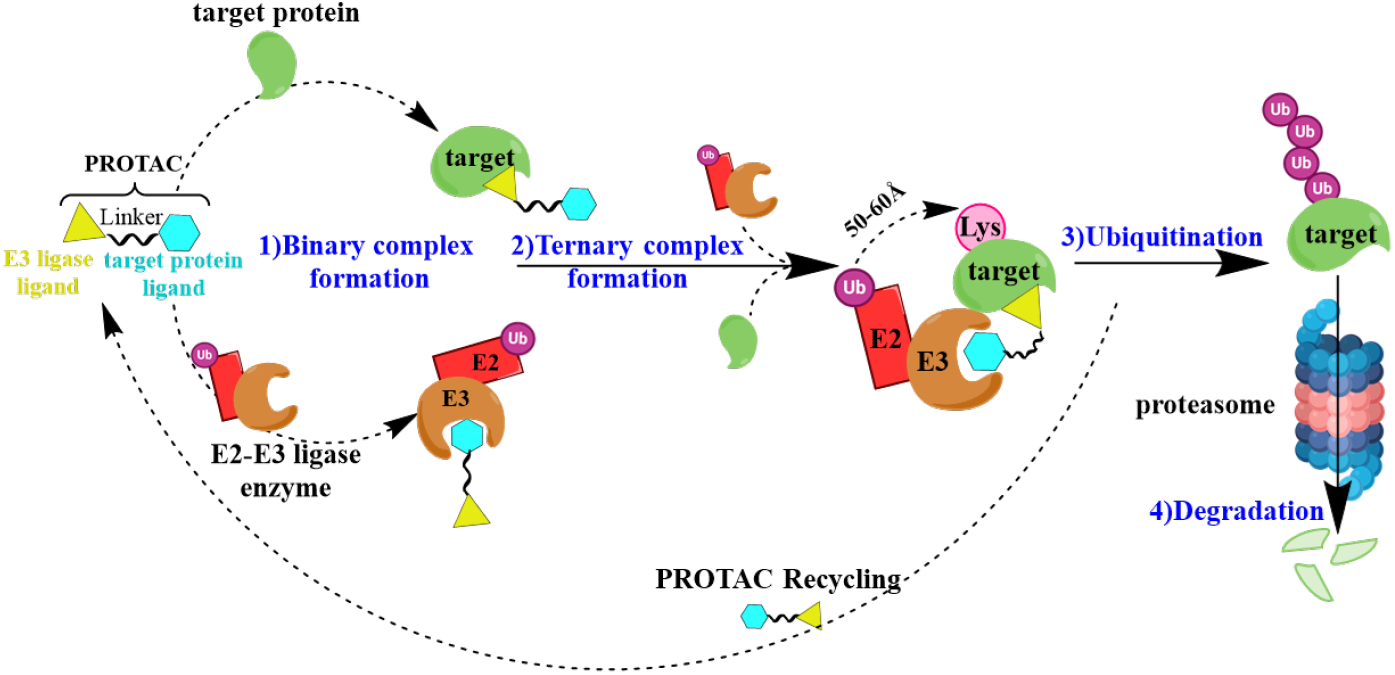
PROTAC-mediated protein degradation. PROTACs are heterobifunctional molecules designed to induce targeted protein degradation. The figure illustrates the mechanism by which PROTACs function: one end of the PROTAC molecule binds to the target protein, while the other end recruits an E3 ligase. This interaction facilitates the ubiquitination of the target protein, marking it for degradation by the proteasome. The process results in the efficient and selective degradation of the target protein, showcasing the potential of PROTACs as powerful tools for targeted therapy.

Traditional research methodologies for PROTACs primarily rely on wet experimental approaches encompassing molecular design, synthesis, biological activity assessments, and clinical trials [1, 2, 7]. While effective, these methods are often time-consuming, labor-intensive, and costly [3, 8]. Recently, machine learning (ML)-based approaches have rapidly progressed in PROTAC research [9]. Classic ML methods typically focus on specific data samples, constructing hybrid descriptors from PROTAC molecules and proteins, and harnessing algorithms to predict or optimize PROTAC properties like degradation capacity [10–13]. Deep learning (DL) methods, benefiting from more complex neural network designs, are capable of deriving latent representations of PROTAC molecules and proteins and capturing their intricate relationships [14, 15]. These approaches encode PROTAC molecules and proteins as inputs, feeding them into deep neural networks such as Recurrent Neural Networks (RNNs) [16–18],

Graph Neural Networks (GNNs) [17, 19–21], or Transformer architectures [21–24], ultimately inferring target outcomes such as generated PROTAC molecules, degradation predictions, facilitating an end-to-end process.

Despite these significant advancements, current DL methods for PROTAC degradation prediction still face notable challenges. *One key challenge is effectively modeling data from POIs, PROTAC molecules, and E3 ligases*. Structural information plays a crucial role in interactions within the PROTAC system involving POIs, PROTAC molecules, and E3 ligases as the interactions are particularly related to substructures in PROTAC molecules and binding sites in the POIs and E3 ligases [4, 25]. PROTACs, as small molecules, inherently possess multi-faceted characteristics, including molecular and atomic properties [26]. However, they are frequently undercharacterized in research, with studies limited to SMILES representations or molecular fingerprints, failing to exploit the multi-scale hierarchical information in PROTAC molecules [13, 17, 22, 23]. Regarding POIs and E3 ligases, many existing methods overlook crucial protein structural information during data modeling. They often rely on raw protein descriptors like amino acid sequences to represent POIs and E3 ligases [13] or sometimes omit the protein component entirely [16, 22–24], which tends to limit the model performance.

*Another major challenge involves explicit learning of substructure interactions within the PROTAC system and the interpretability of computational outcomes*. PROTAC molecules exert their effect by forming ternary complexes with POIs and E3 ligases, primarily through mutual substructure interactions among PROTAC molecules, POIs, and E3 ligases [1, 2]. However, many previous studies have approached this by learning joint features in separate encoders, without explicitly capturing substructure interactions. In these methods, PROTAC, POI, and E3 ligase representations are initially learned separately, with mutual information being implicitly inferred through combination or a black-box module [13, 17, 20, 21], which limits the depth of analysis in PROTAC system interactions. Moreover, the black-box nature of these methods inherently restricts the interpretability of computational outcomes [27]. Without explicit learning of substructure interactions, results become even more challenging to interpret, making it difficult for researchers to comprehend the biological basis underlying the computations.

To address these challenges, we propose *PROTAC-STAN*, a structure-informed deep ternary attention framework for interpretable PROTAC degradation prediction. *PROTAC-STAN* integrates the hierarchical representation of PROTAC molecules, the structural embedding of POIs and E3 ligases, and a ternary attention network to model intermolecular interactions. For the first challenge, we encode hierarchical information in PROTAC molecules with molecular graph, SMILES word frequency mapping, and physicochemical properties across atom, molecule, and property hierarchies, respectively, via a custom Graph Convolutional Network (GCN) [28], retaining critical features. For POIs and E3 ligases, we introduce crucial structural information into protein embedding for protein sequence data without requiring explicit protein structures as input inspired by ESM-S [29, 30], capturing the protein structure information. For the second challenge, we introduce a novel ternary attention network tailored for the PROTAC system. Encoded representations are fed into the ternary attention network to learn substructure interactions among three entities. In this way, we can model interactions among three entities at the atom and amino acid levels, and utilize the learned ternary attention map to visualize the contribution of each substructure to the final result, aiding in revealing their binding mechanisms, thereby enhancing model interpretability. Experiments on the enhanced public PROTAC-DB [31] with refined degradation information demonstrate that *PROTAC-STAN* outperforms state-of-the-art baselines in overall performance, substantially improving degradation prediction accuracy by 10.95%, with AUROC at 0.8833, and F1 score at 0.8588, while enabling significant model interpretability via atomic and residue level visualization of molecule and complex. This computational simulation of the PROTAC system advances PROTAC research, paving the way for future therapeutic development. Our main contributions can be summarized as follows:

- We propose *PROTAC-STAN*, a structure-informed deep ternary attention framework for interpretable PROTAC degradation prediction integrating hierarchical representation of PROTAC molecules, structural embedding of POIs and E3 ligases, and ternary attention network modeling substructure interactions. Our approach effectively models data hierarchically and structurally, retaining critical features and capturing structural information.
- We introduce a novel ternary attention network tailored for the PROTAC system, simulating substructure interactions among POIs, PROTAC molecules, and E3 ligases in nature and yielding interpretable outcomes, offering intuitive insights into PROTAC-mediated protein degradation.
- Experiments on the enhanced PROTAC-DB with refined degradation information show that *PROTAC-STAN* outperforms state-of-the-art baselines, improving overall performance while enhancing interpretability.

## 2 Results

### 2.1 *PROTAC-STAN* framework

The *PROTAC-STAN* framework, depicted in Figure 2, consists of three feature encoder, a ternary attention network, and an MLP classifier. Given a (POI, PROTAC, E3 ligase) triplet, the inputs are first transformed by separate feature encoders: hierarchical encoder for PROTAC, and structural encoders for POI and E3 ligase. The hierarchical encoder, as detailed in Section 4.2.2, encodes hierarchical information for PROTAC, and the structural encoder integrates structural information for POI and E3 ligase, as explained in Section 4.2.3. Next, the encoded representations are fed into the ternary attention network to learn substructure interactions among the three entities, as described in Section 4.2.4. This network produces a joint POI-PROTAC-E3 representation, with a ternary attention map that visualizes the contribution of each substructure, enhancing model interpretability. Lastly, an MLP classification layer predicts degradation outcomes from the joint representation. By leveraging this framework, our model offers nuanced insights into PROTAC-mediated protein degradation interactions, yielding more accurate predictions and improved result interpretability.

**Fig 2.**
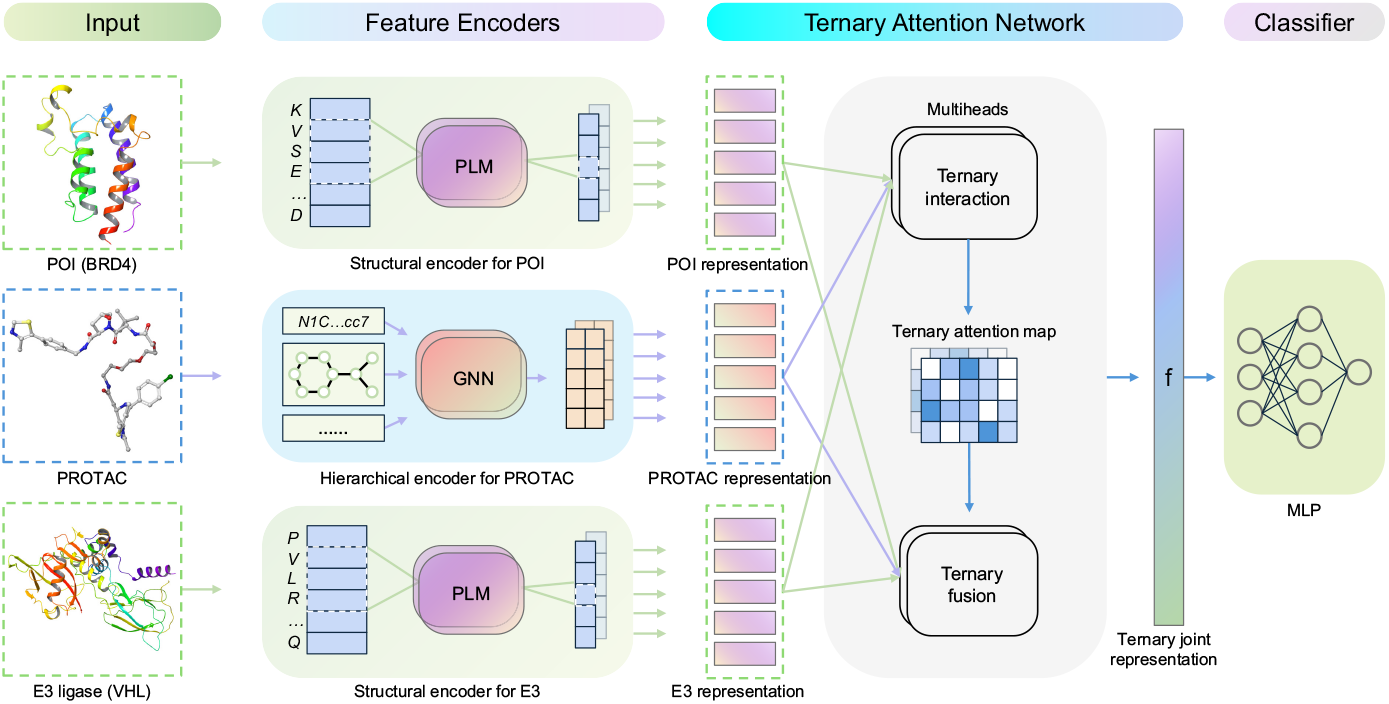
Overview of the *PROTAC-STAN* framework. The input PROTAC molecule, POI (BRD4 example), and E3 ligase (VHL example) are first transformed into hierarchical PROTAC representation and structural POI/E3 representations via feature encoders. The PROTAC encoder encodes hierarchical information across atom, molecule, and property levels, while the structural encoder for POI and E3 ligase incorporates structural information into protein embeddings without needing explicit protein structures. The encoded representations then feed into the ternary attention network to learn substructure interactions. In the first ternary interaction step, the substructure representations of the three entities undergo ternary interaction modeling to generate a ternary attention map that captures interaction intensity. In the second ternary fusion step, these representations are fused into a joint POI-PROTAC-E3 representation **f** over the ternary attention map. Finally, an MLP classifier maps the joint representation to a degradation prediction result.

### 2.2 Performance evaluations

#### 2.2.1 Performance on *PROTAC-fine*

We enrich degradation information to the PROTAC-DB 2.0 [31] as described in 4.1, and construct a refined PROTAC dataset named *PROTAC-fine* which has a total size of 1,503 data samples. We divide *PROTAC-fine* into train/test splits using a random split 0.8/0.2 split, accounting for data leakage by avoiding having the same PROTAC samples in both splits. Data samples where a PROTAC was in the training set are moved out of the test set, leading to 1,202 training samples and 207 samples in the hold-out test set. We trained the *PROTAC-STAN* at the configuration of Table B.1. Here we evaluated the proposed *PROTAC-STAN* with three baselines under random split setting: support vector machine (SVM) [32], random forest (RF) [33], DeepPROTACs [17] using accuracy, AUROC, and F1 score as metrics.

The comparison results are demonstrated in Table 1. Part of the results are adopted from DeepPROTACs because the input POI/E3 ligase data is inaccessible. It is obvious that both the SVM and RF methods show improvement compared to the legacy results from DeepPROTACs, underscoring the data-driven nature of ML approach and affirming our data enrichment efforts in Section 4.1.2. Between the two baselines, SVM performs slightly better than RF, and Morgan [34] fingerprints feature extraction proves marginally more effective than MACCS [35] fingerprints, which is consistent with findings from DeepPROTACs. DeepPROTACs are no better than SVM and RF in accuracy suggesting that classic ML models may perform better than the DL method on raw data when limited data is refined. However, we can see that DL methods perform better overall, especially in AUROC and F1 score, confirming that DL approaches are more adept at modeling complex data connections and are not easily biased in a single metric. Overall, the results demonstrate that *PROTAC-STAN* consistently outperforms the other methods, achieving superior accuracy, AUROC, and F1 score in the public PROTAC dataset. Our data enrichment strategy enhances the model’s ability to perform a more accurate and fitting prediction task, and the design of *PROTAC-STAN* effectively captures hierarchical and structural information from refined data. Furthermore, ternary attention modeling of substructure interactions contributes substantially to the improvements in results. Specifically, *PROTAC-STAN* surpassed the previous sota model, DeepPROTACs, by 10.95% in accuracy, with the highest AUROC at 0.8833, and also exhibited the highest F1 score at 0.8588. This highlights the superior performance of *PROTAC-STAN* compared to previous methods, while also demonstrating its effectiveness in predicting PROTAC degradation.

**Table 1.** Evaluation results of *PROTAC-STAN* and baselines on test set considering data leakage. SVM and RF methods are implemented following DeepPROTACs [17]. * denotes legacy results from DeepPROTACs. **Best** baseline, Second Best baseline, **Shadowed**: Ours.

| Method | Fingerprints | Accuracy | AUROC | F1 score |
| --- | --- | --- | --- | --- |
| SVM [32] | MACCS [35] | 79.40% | 0.7798 | <u>0.7355</u> |
|  | Morgan [34] | **80.40%** | 0.7941 | **0.7547** |
| SVM* [17] | MACCS [35] | 71.48% | 0.7385 | - |
|  | Morgan [34] | 76.76% | <u>0.8196</u> | - |
| RF [33] | MACCS [35] | 76.88% | 0.7431 | 0.6806 |
|  | Morgan [34] | 79.90% | 0.7666 | 0.7059 |
| RF* [17] | MACCS [35] | 68.31% | 0.7973 | - |
|  | Morgan [34] | 69.37% | 0.8117 | - |
| DeepPROTACs* [17] | - | 77.46% | **0.8531** | - |
| *PROTAC-STAN* | - | **88.41%** | **0.8833** | **0.8588** |
| *PROTAC-STAN<sub>pair</sub>* | - | **85.02%** | **0.8571** | **0.8268** |

Notably, *PROTAC-STAN*_*pair*_ is a variant that focuses on pairwise interaction, i.e. the PROTAC and POI pair, the PROTAC and E3 ligase pair, rather than considering all three entities simultaneously of *PROTAC-STAN*. The performance of this variant was worse than that of *PROTAC-STAN*, suggesting that the interaction between proteins may play a crucial role in PROTAC-mediated degradation, a factor that existing methods had not accounted for. This is a fascinating finding that also aligns with and supports recent research [36–38]. Furthermore, we demonstrate the interpretability of our *PROTAC-STAN* method with ternary attention visualization in Section 2.3.

#### 2.2.2 Performance on PROTAC-DB 3.0

Recently, PROTAC-DB 3.0 [39] was released, showcasing a great increase in data volume compared to version 2.0. The total number of entries in the PROTAC table has reached 9,380. However, the entries containing both explicit DC_50_ and D_*max*_ values remain relatively limited, with only 909 entries. Using the same data processing and enrichment methods in Section 4.1 applied to PROTAC-DB 2.0, we compiled a dataset of 3,182 available entries labeled with degradation labels. We filtered out data that appeared in PROTAC 2.0, resulting in a dataset of 1,653 entries. It’s important to note that our model had not encountered any of this data before, and the set is significantly larger than our previous model’s test set, which only contained 200 entries (a 7-fold increase), posing a substantial challenge to the model. We considered two strategies to evaluate the performance of *PROTAC-STAN* on PROTAC-DB 3.0. The first involves randomly sampling a test set of the same size in PROTAC-DB 2.0 from the filtered set and conducting an out-of-distribution (OOD) test. The second strategy is to split the filtered dataset into a fine-tuning set and a test set. Following the same data partitioning approach as in version 2.0, while also accounting for data leakage, we split the filtered entries into 1,322 for fine-tuning and 331 for testing.

Using the same hyperparameter configuration as *PROTAC-STAN*, we conducted OOD test and fine-tune test experiments with our method, SVM, and RF methods. The input POI/E3 ligase data of DeepPROTACs is inaccessible, so we did not consider it. The evaluation results are presented in Figure 5b. As shown, *PROTAC-STAN* outperforms other methods in both tests, achieving the best overall performance. In the left of Figure 5b, our model is the only one to exceed 70% accuracy and consistently leads in other metrics. This is remarkable because none of the test data samples were seen by our model beforehand, yet it performed relatively well on the OOD data, highlighting its robustness. The right of Figure 5b highlights that after fine-tuning, our model gets significant improvements in accuracy, so did SVM and RF, suggesting that a more extensive PROTAC dataset could provide greater support for future PROTAC prediction tasks. We have observed that SVM improves a lot in accuracy, potentially due to its sensitivity to specific data features. SVMs and RF depend on particular features for categorization and face challenges with OOD data as shown in the left of Figure 5b, after fine-tuning on fresh data, their performance experiences improvement. However, their AUROC and F1 scores remained far lower than ours, confirming the resilience of our approach and greater flexibility and advantage of our model over other models in handling new data samples. This further underscores the overall capability of our method. Given the constraints imposed by limited training data in scale, there remains potential for enhancing our model’s performance when applied to novel datasets. Notwithstanding these challenges, incorporating hierarchical and structurally encoded information alongside the utilization of ternary attention mechanisms has enabled our model to attain satisfactory results. This endeavor represents a promising exploration.

**Fig 3.**
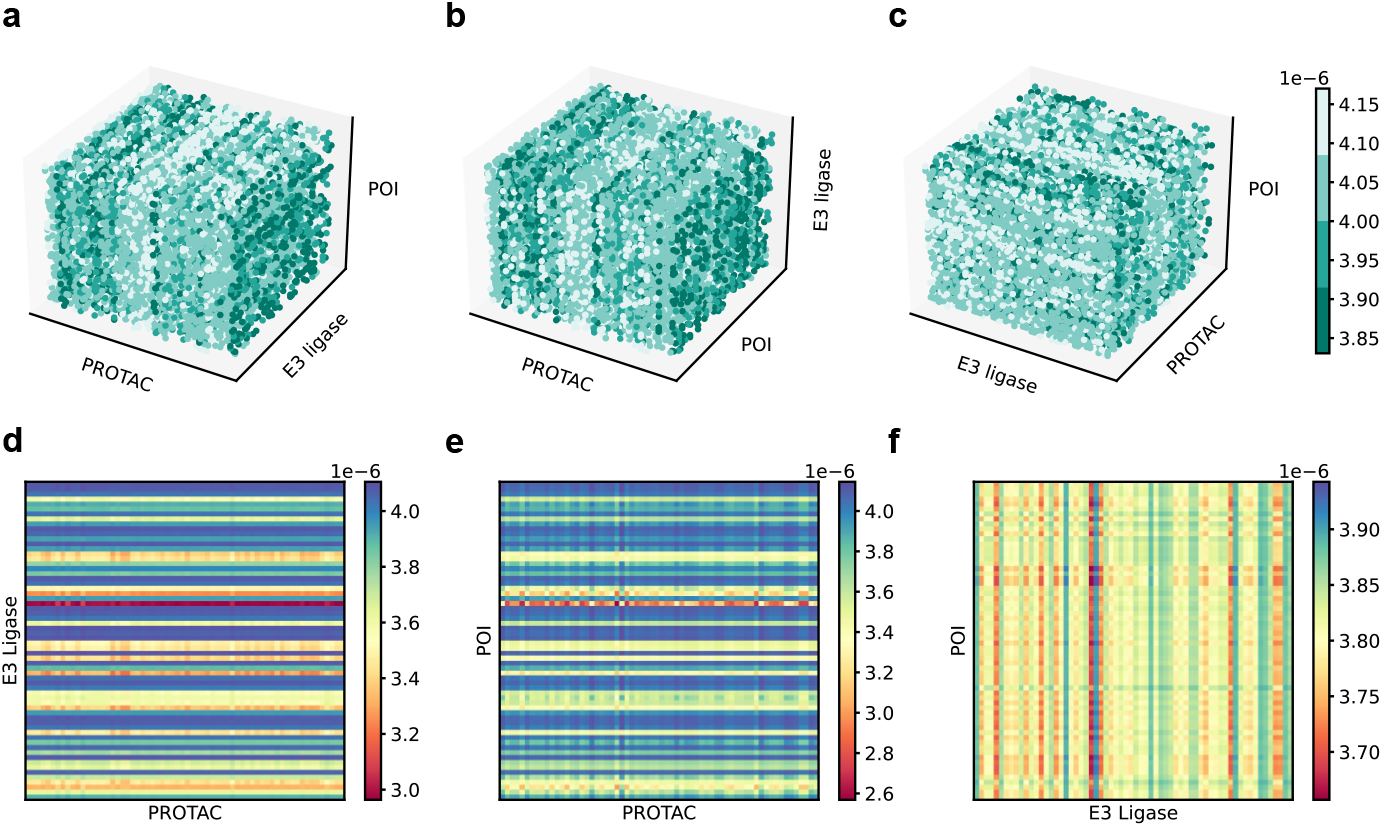
3D attention visualization in multi-view and pairwise attention map visualization. **a**, Front view. Observation from front to back of attention map. Strong bar patterns of PROTAC interacting with POI. **b**, Top view. Observation from top to bottom of attention map. Strong bar patterns of PROTAC interacting with E3 ligase. **c**, Side view. Observation from left to right of attention map. Weak patterns between POI and E3 ligase. **d**, The PROTAC-E3 ligase pairwise attention map. **e**, The PROTAC-POI pairwise attention map. **f**, The E3 ligase-POI pairwise attention map.

**Fig 4.**
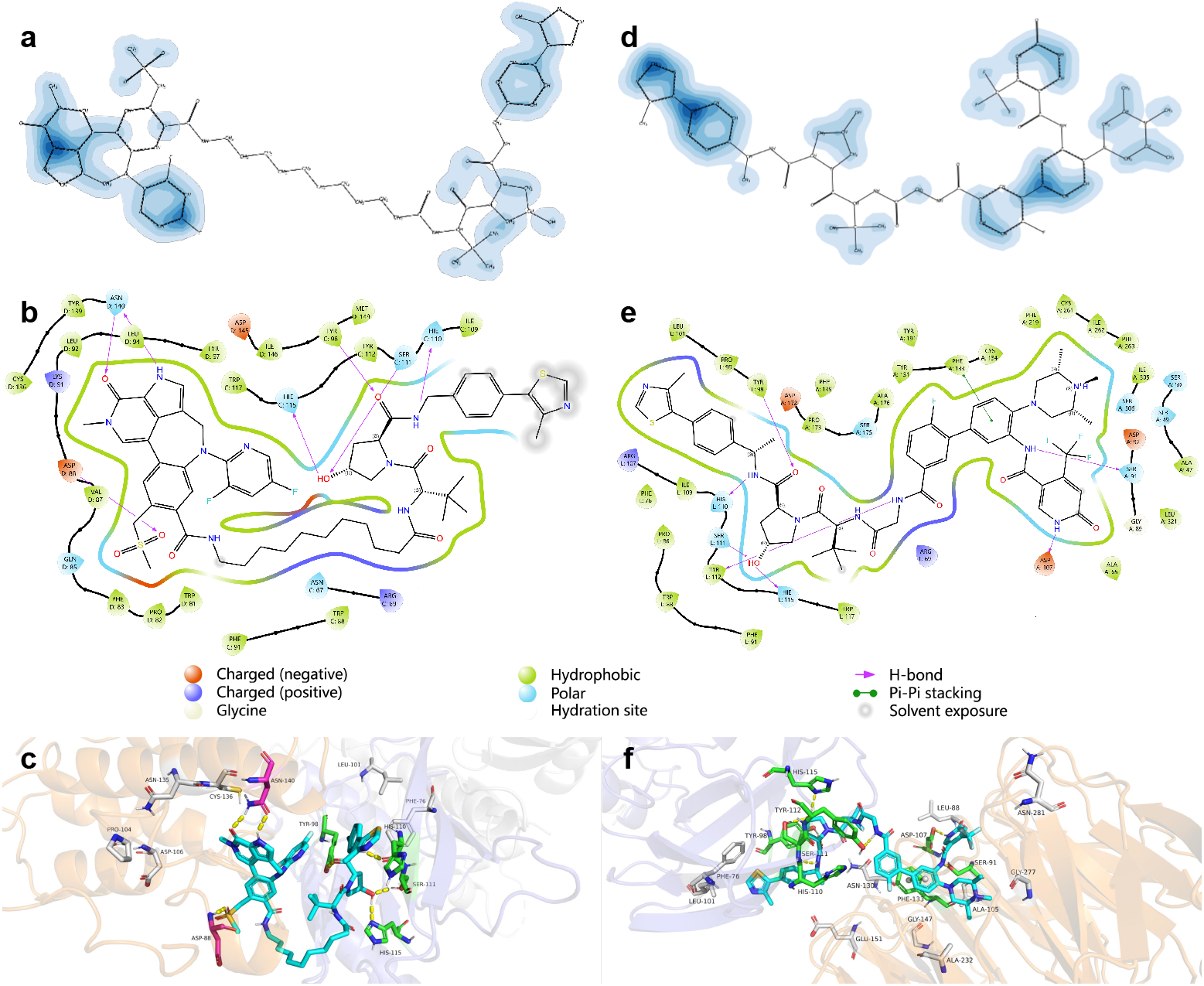
Ternary attention visualization on the PROTAC molecule and 3D Complex. Left (a-c) is from PDB 7KHH and right (d-f) from PDB 7JTP. **a, d**, 2D PROTAC molecule visualization with weighted atoms. The deeper the color, the higher the weight of the atom. **b, e**, 2D interaction visualization extracted from the complex with legend below.**c, f**, 3D pocket visualization with key residues. Top weighted residues are marked in gray, the actual interacting residues in green, and overlapped residues in magenta.

**Fig 5.**
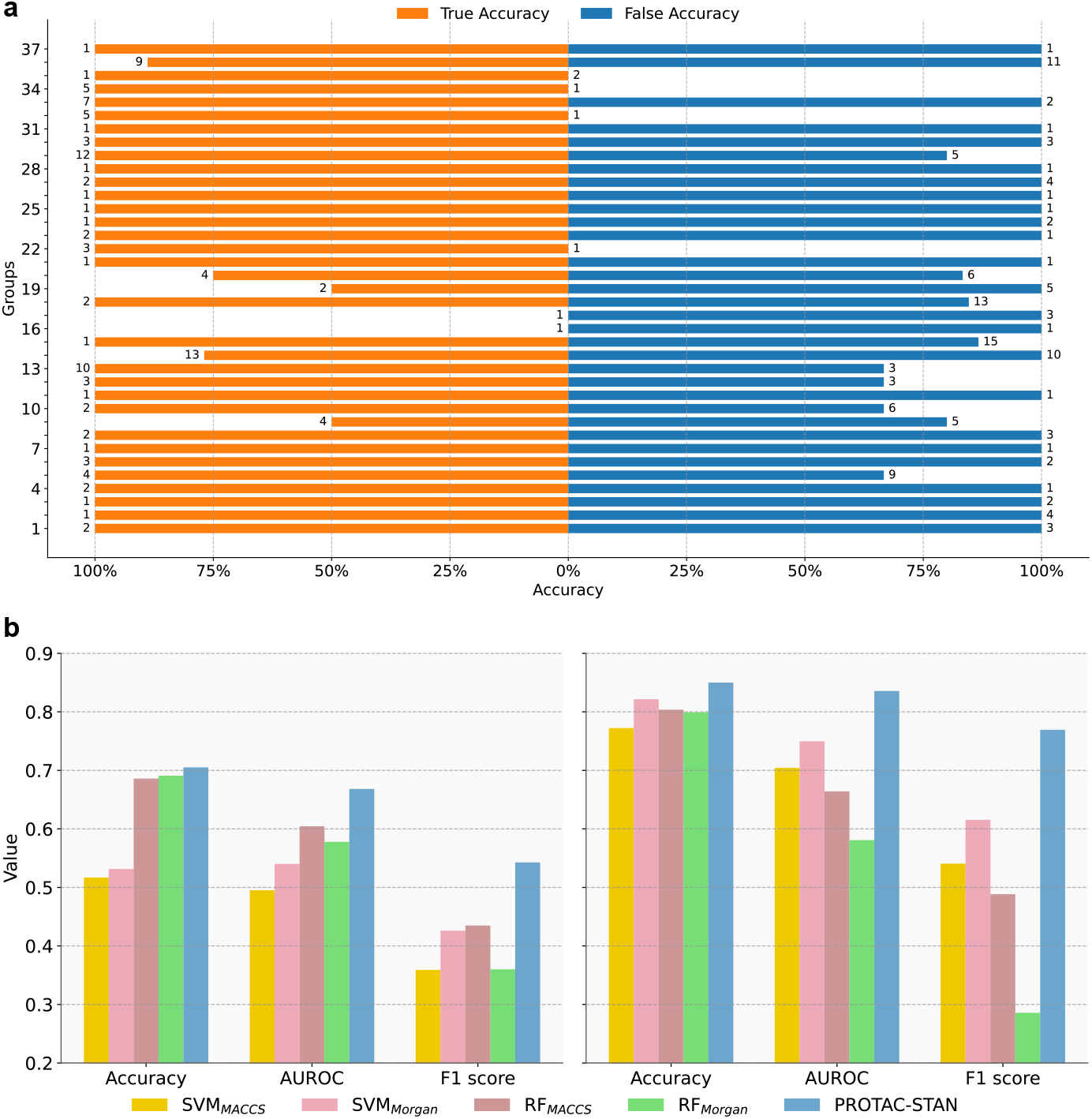
a, Linker sensitivity evaluation through grouped degradation prediction. Samples in each group have fixed E3 ligase-E3 ligand and POI-warhead pair, and varied linker. The number of both ends in figure are sample numbers in each group. **b, Evaluation on the dataset constructed from the most recent PROATC-DB 3.0**. Left is out-of-distribution test and right is fine-tune test.

### 2.3 Interpretability with ternary attention visualization

#### 2.3.1 Interpreting in 3D attention map

The strength of *PROTAC-STAN* is to utilize the ternary attention map to provide interpretability and insights for understanding how each substructure in three entities contributes to the PROTAC degradation prediction results. We take one sample 8 (Compound ID: 22) from PROTAC-DB as an example and retrieve the generated three-dimensional attention map after the ternary interaction process. We visually represent it in 3D across multiple perspectives, as shown in Figure 3a-c. The shallow green hue signifies intense interaction, medium green denotes moderate interaction, and deep green indicates minimal or no interaction. In Figure 3a, the front view reveals discernible patterns in the PROTAC dimension’s relationship to the POI dimension, suggesting specific interplays that significantly contribute to the formation of the ternary complex. In particular, we can observe that the feature images display vertical bars of varying shades. These distinct shades correspond to the interactions between specific locations on the PROTAC molecule and targeted regions on the POI during the learning process facilitated by ternary attention. Similarly, this pattern is evident in the top view in Figure 3b, further underscoring the importance of these interactions. Notably, the side view in Figure 3c exhibits weak patterns between POI and E3 ligase aligns with the detailed pairwise attention relationships presented in Section 2.3.2 indicating the comparatively weak interaction between POI and E3 ligase. Through direct 3D visualization of the ternary attention map, we can gain a macroscopic understanding of the potential interplay patterns among PROTAC, E3 ligase, and POI, which may facilitate our comprehension of their ternary complex formation process.

#### 2.3.2 Interpreting in 2D attention map

The three-dimensional attention map quantifies the interactions among PROTAC, E3 ligase, and POI, allowing for the extraction of pairwise relationships in a straightforward manner. By employing a mean-based approach along one dimension, we derive the other two pairwise attention maps, yielding the PROTAC-E3 ligase attention map, the PROTAC-POI attention map, and the E3 ligase-POI attention map, as depicted in Figure 3d-f. The visualizations reveal that the attention values are notably elevated for PROTAC-E3 ligase and PROTAC-POI interactions, whereas relatively diminished for E3 ligase-POI interactions. These intense values suggest the presence of specific regions in E3 ligase and POI, closely interacting with the PROTAC molecule. The medium E3 ligase-POI interaction observed in 3f corroborates findings in Section 2.2.1, which underscored the significance of protein-protein interactions in PROTAC-mediated degradation and their much weaker interaction compared to PROTAC-E3 ligase/POI interactions. The 2D visualizations facilitate a deeper understanding of the pairwise relationships within the PROTAC system, providing valuable insights into PROTAC-mediated protein degradation. More examples of attention map visualizations are provided in Appendix C.

#### 2.3.3 Interpreting in molecule and complex

Beyond visualizing the attention maps, *PROTAC-STAN* ‘s capability can most importantly map these weights back onto the atom level of the PROTAC molecule and amino acid residue level of E3 ligase and POI, enabling interpretable analysis at molecular and complex levels. We mapped ternary attention back to the atomic level of PROTAC and to the residue level of E3 ligase and POI, and performed visualization on PROTAC molecules and 3D complexes. The visualized results are shown in Figure 4, along with 2D interaction diagrams extracted from 3D complexes. Two examples (PDB ID: 7KHH [40] and 7JTP [41]) were used to demonstrate this, where we obtained their 3D crystal structures from Protein Data Bank (PDB) [42], prepared the proteins using Maestro, removed excess solvents, and filled in missing loops to obtain optimized structures.

We extracted the atomic weights of PROTAC from ternary attention maps and colored 60% of atoms, as shown in Figure 4a and 4d. The deeper the color, the higher the weight of the atom. We used Maestro to produce 2D interaction diagrams among PROTAC, E3 ligase, and POI from 3D complexes, with a range set at 3.5 Å, as shown in Figure 4b and Figure 4e. We find the main interactions between PROTACs and proteins are hydrogen bonding and π-π stacking interactions, with the interacting atoms mainly located at both ends of PROTAC, which is consistent with their actual binding modes (PROTAC warhead binds to POI, E3 ligand binds to E3 ligase). Take Figure 4a for example, the 2D molecular visualization of PROTAC accurately identified multiple key interaction regions, as well as corresponding atoms or groups. Specifically, in the left part of the molecule in Figure 4b, the oxygen atom (C=O) from the 2-Pyridone ring interacts with residue ASN D:140, and the NH group from the adjacent pyrrole ring also interacts with ASN D:140, forming stable hydrogen bonds. The oxygen atom (S=O) from the methylsulfonyl group interacts with ASP D:88, forming a hydrogen bond. In the right side of the molecule in Figure 4b, the hydroxyl group (OH) from the 3-hydroxypyrrole ring interacts with residue SER C:111 and HIE C:115, forming hydrogen bonds. Additionally, the carbonyl oxygen atom (C=O) from the amide group interacts with TYR C:98, and the nitrogen atom (NH) from the amine group interacts with HIE C:110, also forming hydrogen bonds. These visualizations reflect the detailed molecular interactions between PROTACs and protein binding pockets, intuitively enhancing our understanding of how PROTAC forms ternary complexes with E3 ligase and POI.

Figure 4c and Figure 4f show the binding situation within 3D pockets. We used PyMOL to implement this, with a range set at 3.5 Å. From ternary attention maps, we extracted the residue weights of E3 ligase and POI, marked the top important weights in gray, and marked the actual interacting residues in green, overlapped residues in magenta, hydrogen bonds in yellow, and π-π stacking interactions in green. The blue-purple part in background represents E3 ligase, while the orange part represents POI. In Figure 4c, it can be seen that residue ASN-140 and ASP-88 were accurately marked as high-weight residues, which formed hydrogen bonding interactions with PROTAC molecules. Other top-weight residues not forming interactions, such as CYS-136, and PHE-76, are also located near the interacting residues, which is quite impressive and indicates that our method can provide a certain level of interpretability for 3D complexes. In Figure 4f, the interacting residues are relatively concentrated, although there are no overlapping residues. However, our high-weight residues such as PHE-76, LEU88, ASN-130, GLY-147, and ALA-105 are all very close to the interacting residues, which suggests that our ternary attention learning has a certain trend perception ability for key residues and can potentially help us understand PROTAC forming ternary complexes.

### 2.4 Linker sensitivity of *PROTAC-STAN*

The linker in PROTAC plays a pivotal role in determining its biodegradative efficacy [43], serving as a critical component that influences many aspects of the design and function of PROTAC. In this section, we investigate the sensitivity of our method to linker variations. We aligned the PROTAC samples in our refined dataset with additional data table containing warheads, linkers, and E3 ligands [31], resulting in a total of 1,323 data entries, which include (PROTAC, warhead, linker, E3 ligand) matches. We then grouped these entries by (warhead, E3 ligand) pairs and discarded groups containing only true or false samples, yielding 251 data entries across 37 groups. We evaluated all samples using *PROTAC-STAN*, achieving a degradation prediction accuracy of 88.45%, showing our method’s excellence. We then conducted experiments for each group, and the results are presented in Figure 5a. Among the 37 groups, accurate predictions for both true and false samples were observed in 20 groups, with overall accuracy being relatively balanced between true and false predictions. We observed that groups exhibiting 0% prediction accuracy primarily consist of single samples, which highlights the urgent need for more PROTAC data to enhance our current method. In each group, the E3 ligase-E3 ligand and POI-warhead remained constant while the linker varied. Our method effectively predicted the correct degradation outcomes, indicating that *PROTAC-STAN* exhibits sensitivity to linker variations during degradation predictions. This finding enhances the applicability of *PROTAC-STAN*, enabling its future integration with linker design methodologies for the discovery of new PROTAC candidates.

### 2.5 Ablation study

#### 2.5.1 Ablating components of *PROTAC-STAN*

Here we perform ablation experiments to evaluate the key components of the *PROTAC-STAN* framework introduced in Section 2.1. To this end, we use a series of ablation schemes. The Raw scheme replaces each component in *PROTAC-STAN* with the PROTAC SMILESNet encoder, the POI/E3 ligase Ngrams encoder, and concatenation fusion for comparison. We then analyze the impact of each component individually. Specifically, the -S1 scheme isolates the effect of the PROTAC hierarchical encoder, the -S2 scheme isolates the effect of the POI/E3 ligase structural encoder, and the -S scheme combines -S1 and -S2 to assess the overall effect of feature encoding. Additionally, the -T scheme evaluates the influence of the ternary attention network. Lastly, we compare the overall performance enhancement of *PROTAC-STAN* against the Raw scheme. From Table 2, it is evident that the Raw model constructed from baselines performs adequately in predicting degradation, thus validating previous work. The -S model improves the F1 score from 0.7738 to 0.8333, highlighting the effectiveness of our structure-informed feature encoders, with -S1 more contributing than -S2. The -T model is comparable to Raw due to the simple encoding of Raw; however, the notable difference lies in that -T introduces a ternary attention network to fuse POI, PROTAC, and E3 ligase as well as modeling ternary interactions among them. Moreover, incorporating -S and -T components, i.e. *PROTAC-STAN*, results in a 6.77% improvement in accuracy and achieves 0.8833 in AUROC and 0.8588 in F1 score, thereby validating the superior performance of our method.

**Table 2.** Ablation Study on *PROTAC-STAN* Components. The Raw scheme is a base constructed from baselines, the -S1 and -S2 schemes ablate the PROTAC hierarchical and POI/E3 ligase structural encoders. The -S scheme combines -S1 and -S2 to assess feature encoding effects. The -T scheme evaluates the ternary attention network over concatenation. **Best**, Second Best.

| Component | Raw | -S1 | -S2 | -S | -T | -STAN |
| --- | --- | --- | --- | --- | --- | --- |
| SMILESNet encoder [17] | ✓ |  | ✓ |  | ✓ |  |
| Hierarchical encoder |  | ✓ |  | ✓ |  | ✓ |
| Ngrams encoder [13] | ✓ | ✓ |  |  | ✓ |  |
| Structural encoder |  |  | ✓ | ✓ |  | ✓ |
| Concatenation [13, 17] | ✓ | ✓ | ✓ | ✓ |  |  |
| TAN fusion |  |  |  |  | ✓ | ✓ |
| Accuracy | 81.64% | <u>85.99%</u> | 81.64% | 85.51% | 80.68% | **88.41%** |
| AUROC | 0.8109 | <u>0.8671</u> | 0.8229 | 0.8631 | 0.8108 | **0.8833** |
| F1 score | 0.7738 | <u>0.8380</u> | 0.7889 | 0.8333 | 0.7753 | **0.8588** |

#### 2.5.2 Ablating ternary attention network (TAN)

We further investigate the TAN module of *PROTAC-STAN* by examining different fusion methods for POI, PROTAC, and E3 ligase features, varying TAN heads, and input orders. We compare TAN fusion against concatenation fusion [13, 17] and LMF 13 [44], a low-rank multimodal fusion method originally designed for visual, audio, and language inputs. As shown in Appendix Table B3, Concatenation and LMF are comparable to TAN_1_ but perform less well than TAN_2_. This indicates that all three methods are capable of fusing the three features, but in terms of methods, Concatenation is significantly naive compared to LMF and TAN. LMF and TAN are more complex to train, which means they are better performed at the same complexity. Additionally, TAN introduces a ternary attention map, providing better interpretability compared to Concatenation and LMF. For multi-head configurations, we find that multiple TAN heads outperform a single head, but with increased parameters. This is why we opted not to add more heads, considering the parameter growth and dataset volume. Additionally, we analyze the impact of different input permutations for POI, PROTAC, and E3 ligase, revealing that the order of inputs has a minimal effect on final performance. The permutations 123 and 312 (corresponding to PROTAC, E3 ligase, and POI) yield the best results, while other orders show similar performance, indicating that the influence of input feature order is minor.

## 3 Discussion

In this study, we introduce *PROTAC-STAN*, an end-to-end structure-informed deep ternary attention framework designed for interpretable PROTAC degradation prediction. Our approach integrates hierarchical representations of PROTAC molecules, structural embeddings of POIs and E3 ligases, and a ternary attention network to model substructure interactions effectively. By leveraging hierarchical and structural data modeling, *PROTAC-STAN* retains essential features and captures critical structural information. The novel ternary attention network, specifically tailored for the PROTAC system, simulates substructure interactions among POIs, PROTAC molecules, and E3 ligases, yielding interpretable outcomes and providing intuitive insights into PROTAC-mediated protein degradation. Experiments on the refined dataset from PROTAC-DB demonstrate that *PROTAC-STAN* outperforms state-of-the-art deep learning models and traditional machine learning approaches in predicting PROTAC degradation, imporving accuracy by 10.95%, with AUROC at 0.8833, and F1 score at 0.8588, while also facilitating extensive model interpretability via atomic and residue-level visualization of molecules and complexes. The architecture of *PROTAC-STAN* facilitates an in-depth understanding of how PROTACs interact with proteins and ligases, moving beyond traditional models that only consider binary interactions. This comprehensive approach enables the model to capture complex interaction dynamics, contributing to the advancement of PROTAC drug discovery and offering insights into optimizing drug design for enhancing efficacy and selectivity. In conclusion, *PROTAC-STAN* has achieved notable success in surpassing other similar algorithms. We anticipate that, with the future expansion of PROTAC datasets, this model will demonstrate enhanced robustness. Moreover, the incorporation of a broader range of information, including dynamic factors such as binding kinetics, is expected to further improve the model’s performance.

## 4 Methods

### 4.1 Data preparation

#### 4.1.1 Dataset

We sourced the original data from PROTAC-DB 2.0 [31], which is structured into four main tables detailing information on PROTACs, Warheads, Linkers, and E3 Ligands. The database includes chemical structures, biological activities, and physicochemical properties extracted from experimental literature. The primary table for PROTACs comprises 5388 entries, representing 3270 unique PROTAC molecules. Duplicate entries typically denote the same molecule tested under different experimental conditions. Each entry includes the PROTAC’s SMILES, the POI and E3 ligase’s UniProt ID. Crucially, the dataset contains information on PROTAC degradation activity, including DC_50_ (half-maximal degradation concentration) and D_*max*_ (maximum level of protein degradation). These metrics are standard indicators of a PROTAC’s capability to degrade its target protein [45], with lower DC_50_ and higher D_*max*_ values suggesting higher degradation activity.

#### 4.1.2 Data Enrichment

Due to a significant proportion of PROTACs in the original database lacking DC_50_ and D_*max*_ data, the availability of degradation-informed data is notably limited, posing challenges for deep learning applications. Among the 5,388 entries cataloged, merely 362 possess detailed records of both DC_50_ and D_*max*_ values. Our in-depth analysis of the database in Appendix B.3 indicates that it is possible to derive additional degradation activity information from experimental descriptions related to *Assay (*DC_50_*/*D_*max*_*), Percent degradation*, and *Assay (Percent Degradation)*. Specifically, for each entry, we first check for explicit DC_50_ or D_*max*_ values; if none are present, we then extract implied DC_50_ and D_*max*_ values from available experimental descriptions. This effort has enabled us to preliminarily yield 1631 entries, effectively increasing the original data volume by 350%.

#### 4.1.3 Degradation Labeling

The quantification of a PROTAC’s ability to degrade target protein is commonly assessed using DC_50_ and D_*max*_ [45]. To effectively train our deep learning model, we established a robust labeling protocol based on that used in DeepPROTACs [17] for processing degradation labels. This protocol considers both explicit data points, where DC_50_ and D_*max*_ values are directly available, and implicit scenarios where these values need to be inferred from experimental observations. This tiered protocol allows us to effectively utilize available data and train our model for accurate PROTAC degradation prediction. The specific criteria are as follows:

- **Explicit** DC_50_ **and** D_*max*_ **values**: A PROTAC is labeled as high degradation activity if DC_50_ is less than 100 nM or D_*max*_ reaches or exceeds 80%. Conversely, it is considered to have low degradation activity if DC_50_ is greater than or equal to 100 nM and D_*max*_ is below 80%.
- **Implicit** DC_50_ **and** D_*max*_ **values**: When these values are not directly obtained, the PROTAC’s degradation activity is inferred. If the maximum DC_50_ dosage is under 100 nM, the PROTAC is deemed to have high activity. If not, but the D_*max*_ at DC_50_ = 100 nM reaches or exceeds 80%, it is still considered highly active. All other instances suggest low activity.

Following these labeling criteria, we identified 620 entries as possessing high degradation activity and 1,011 entries as low. We name the refined dataset PROTAC-fine and make it publicly available. These enhancements to PROTAC-DB significantly bolster the database’s comprehensiveness, offering substantial support for future research and applications in the field.

### 4.2 *PROTAC-STAN* Architecture

#### 4.2.1 Problem formulation

In PROTAC degradation prediction, the task is to determine whether a triplet of a POI, a PROTAC molecule, and an E3 ligase will form a ternary complex to degrade the POI via the UPS. Given a POI 𝒫_*p*_, a PROTAC molecule 𝒢, and an E3 ligase 𝒫_*e*_, PROTAC degradation prediction aims to learn a model ℳ to map the joint feature representation space 𝒫_*p*_ × 𝒢 × 𝒫_*e*_ to a degradation label l ∈ {0, 1}, where 1 indicates high degradation and 0 indicates low degradation. Major notations used in this paper are provided in Appendix Table A1. We present the *PROTAC-STAN* architecture step by step, and the overview of *PROTAC-STAN* is illustrated in Figure 2.

#### 4.2.2 Hierarchical encoder for PROTAC molecules

The hierarchical molecular feature encoder transforms an input PROTAC molecule into a hierarchical 2D molecular graph 𝒢(⟨𝒱, ℋ⟩,ε) as illustrated in Figure 6a. For node information 𝒱, each atom node is initially defined by its chemical properties using the PyG package. Each atom is characterized by 9 atomic features: atomic number, chirality, degree, formal charge, number of hydrogen atoms, number of radical electrons, hybridization state, aromaticity, and ring membership. Consequently, the node feature for each graph is denoted as 𝒱= {v_1_, v_2_, … }, where v_*i*_ is the i-th node feature, with |𝒱| represents the number of nodes. Regarding edge information, it is represented as *ε* = (*ε*_*e*_, *ε*_*a*_), where *ε*_*e*_ ∈ ℝ^*ϵ×*2^ denotes the edge connections, and *ε*_*a*_ ∈ ℝ^*ϵ×*3^ encodes three bond attributes: bond type, bond stereochemistry, and conjugation status, with ϵ represents the number of edges in the graph.

**Fig 6.**
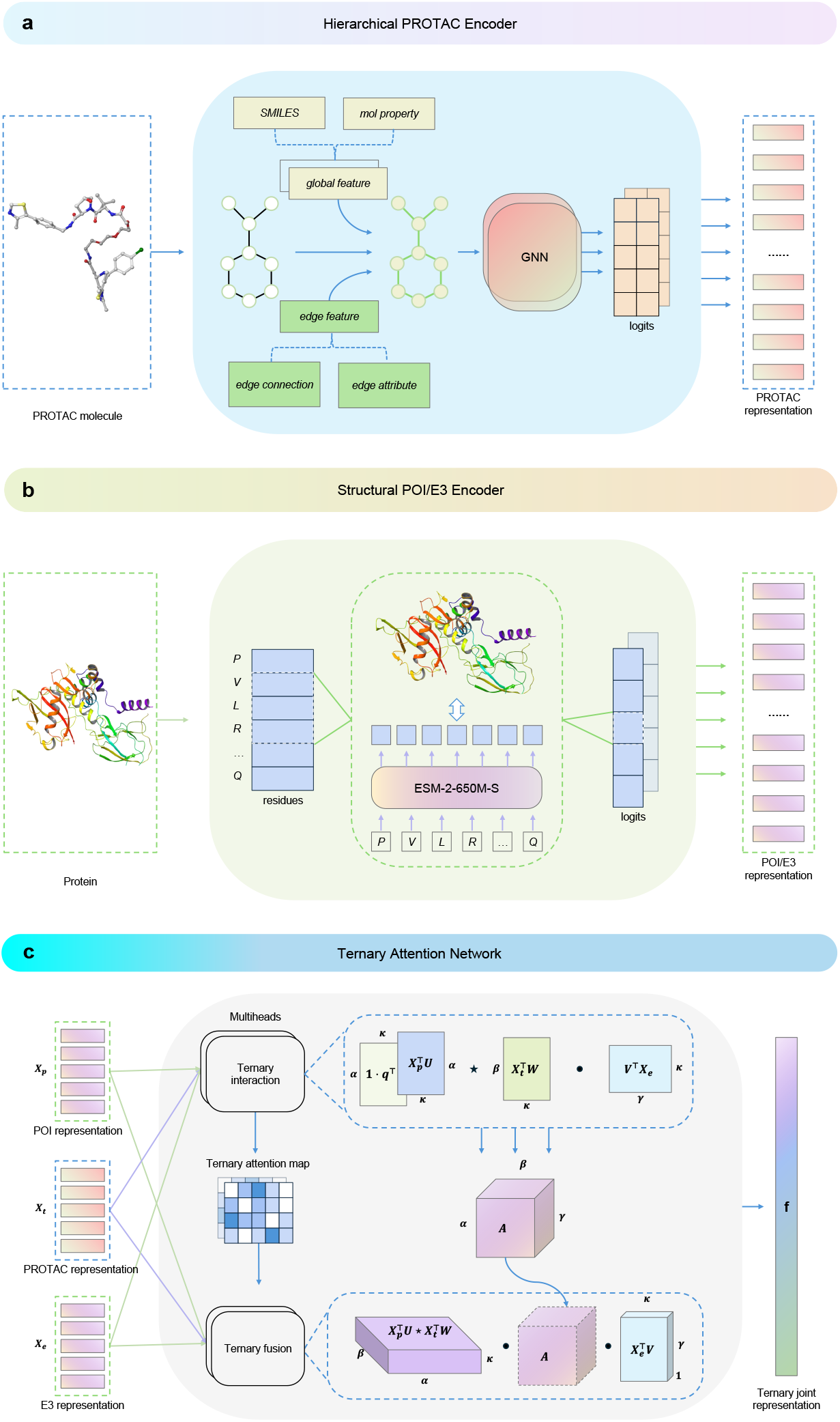
a, Hierarchical PROTAC encoder. Given a PROTAC molecule, the encoder represents it across atom, molecule, and property hierarchies. **b, Structural protein encoder**. Given a POI or E3 ligase, the encoder transforms its amino acid sequence into a structure-informed representation. **c, Ternary attention network**. Given the POI, PROTAC and E3 representation **X**_*p*_, **X**_*t*_ and **X**_*e*_, a ternary attention map **A** and joint POI-PROTAC-E3 representation **f** are obtained. ⋆ denotes the Einstein summation convention *ik, jk → ijk*, and *·* denotes the matrix multiplication.

To represent hierarchical information in 𝒢, we construct a global feature ℋ appended to each atom node feature as ⟨𝒱, ℋ⟩, = {Concat(v_*i*_, ℋ) | v_*i*_ ∈ 𝒱} using SMILES encoding and molecule properties. This hierarchical representation allows us to capture both local atomic information and global molecular characteristics. We first map the PROTAC SMILES into frequency encoding 𝒮, with a maximum allowed string length |𝒮| of from Appendix B.4 using the lead-like molecules encoding table which captures the statistical chemical composition and structural motifs present in the molecule. This table collected by Li et al [17] encodes the 39 most frequent characters from 1 to 39, based on counts from the ZINC database [46]. Nine molecule properties are directly obtained from the PROTAC table in PROTAC-DB [31], denoted as a real-valued vector ℛ. To ensure consistent scaling across diverse properties, these values are standardized using mean and standard deviation normalization, detailed in Appendix B.5. Thus, we construct the global feature ℋ = Concat (𝒮, ℛ), where ℋ ∈ ℝ^|𝒮|+9^, and, 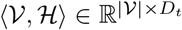, where D_*t*_ = 9 + |𝒮 | + 9, leading to the input PROTAC feature denoted as 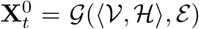. The hierarchical encoding enables our model to learn both detailed atomic interactions and the broader influence of molecular properties on PROTAC degradation activity.

Finally, the hierarchical graph representation of PROTAC molecules was learned through a two-layer GCN module. GCNs extend convolutional operations to non-Euclidean domains, enabling the processing of irregular graph structures. In this process, atom feature matrices are updated by aggregating information from adjacent atoms and their associated chemical bonds. This propagation mechanism inherently captures the substructure information of PROTAC molecules, preserving essential features for subsequent explicit learning of substructure interactions with POI and E3 ligase. The hierarchical PROTAC encoder can be formally expressed as:

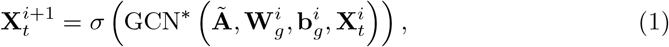

where GCN^*^(·) is our modified GCN to adopt edge attributes, 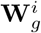 and 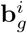 are he GCN’s layer-specific learnable weight matrix and bias vector; **Ã** is the adja-cency matrix with added self-loops in molecular graph 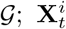 is the i-th hidden representation; and σ(·) denotes a non-linear activation function.

#### 4.2.3 Structural encoders for POIs and E3 liagses

Due to the scarcity of crystal structures for PROTAC-mediated ternary complexes, it is challenging to identify the accurate binding pocket of POI and E3 ligase. Moreover, the sequence data for POIs and E3 ligases is readily available. Therefore, we leverage the abundance of protein sequence data and consider the entire protein structure. As illustrated in Figure 6b, the structural protein encoder consists of a protein language model (PLM) embedder and a linear adapter, which introduces crucial structural information into protein embedding for protein sequence data without requiring explicit protein structures as input. This approach, inspired by ESM-S [29, 30], allows us to represent proteins structurally without relying on explicit 3D structures. The structural encoder offers a powerful alternative when direct structural data is limited, enabling our model to learn and utilize the inherent structural information encoded within protein sequences.

Specifically, the original data for POI and E3 ligase derived from PROTAC-DB are their UniProt IDs [47]. We first retrieve the protein amino acid sequence 𝒫_*p*_ = [r_*p*1_, r_*p*2_, …] for POI and 𝒫_*e*_ = [r_*e*1_, r_*e*2_, …] for E3 ligase, where each r_*i*_ corresponds to the type of the i-th residue and Appendix B.6 provides the length distribution of all proteins. For each protein of POIs and E3 ligases, we construct a residue protein view and then truncate the protein to a maximum allowed length of 1022 [29] using the TorchDrug package. The PLM embedder ϕ is built upon the ESM-S [30] model, which uses remote homology detection to distill structural information into ESM [29], with ϕ_*p*_ for POI and ϕ_*e*_ for E3 ligase. In the PLM embedder module, the protein sequence data is processed into the token representation via several sel-attention and feed-forward networks, and then the sequence representation of a fixed length θ_**p**_ = 1280 is generated in an averaging way. Finally, The input POI and E3 ligase data are denoted as 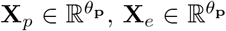, respectively. The structural POI and E3 ligase encoders are described as follows:

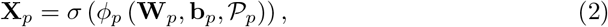

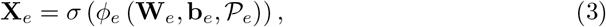

where **W**_*p*_, **W**_*e*_ and **b**_*p*_, **b**_*e*_ are the learnable weight matrices (adopters) and bias vectors after the PLM embedder module. **X**_*p*_ is the structural embedding of the POI and **X**_*e*_ is the structural embedding of the E3 ligase.

#### 4.2.4 Ternary attention network for interaction learning

To capture the complex interplay among POIs, PROTAC molecules, and E3 ligases, we introduce a novel ternary attention network (TAN) tailored for the PROTAC system. Traditional attention [48–50] mechanisms often focus on pairwise interactions, but PROTACs inherently involve three distinct entities. Our TAN, illustrated in Figure 6c, extends this concept to model the intricate relationships among all three simultaneously through a ternary attention map that quantifies interaction intensities and a fusion layer that integrates these interactions into a unified representation. We generalize previous Bilinear Attention Networks (BAN) [49] which are designed for Visual Question Answering (VQA) to a unified network. Interestingly, BAN can be viewed as a subset of our TAN method, as discussed in Appendix A.2, validating the integrity and enhancing the generality of our approach.

TAN comprises two essential components: **(i) a ternary attention map to capture interaction intensity, and (ii) a ternary fusion layer over the attention map to extract joint POI-PROTAC-E3 representation**. TAN enables simultaneous modeling of interactions among all three entities, yielding comprehensive interaction data. This network is particularly well-suited for PROTAC system modeling, as it facilitates concurrent learning of substructure interactions, a crucial aspect in understanding PROTAC-mediated protein degradation. This simultaneous modeling of multi-entity interactions allows our model to learn intricate substructure relationships, leading to improved prediction accuracy and greater interpretability of PROTAC-mediated protein degradation.

Given the learned hidden representation of POI, PROTAC molecule and E3 ligase 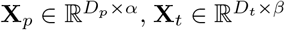, and 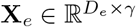 after separate hierarchical molecular and structural protein encoders, where 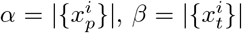, and 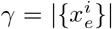 denote the number of encoded substructures in POI, PROTAC molecule, and E3 ligase, respectively. Their ternary interaction can obtain a three-dimension ternary attention map A ∈ ℝ^*α×β×γ*^ :

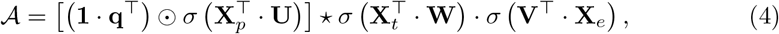

where 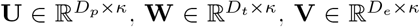 represent learnable weight matrices for POI, PROTAC, and E3 ligase representations, respectively. **q** ∈ ℝ^*κ*^ denotes a learnable weight vector, **1** ∈ ℝ^*α*^ is a constant vector of ones. ⊙ represents the Hadamard (element-wise) product, ⋆ signifies the Einstein summation convention (einsum) [51], specially ik, jk → ijk, and · denotes standard matrix multiplication. The elements of 𝒜 quantify the interaction intensity among (POI, PROTAC, E3) substructural triplets, correlating with potential binding sites and molecular substructures. For a more intuitive understanding of the ternary interaction, an individual element 𝒜_*i,j,k*_ from Equation 4 can be expressed as:

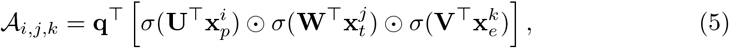

where 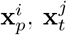, and 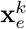 represent the i-th, j-th, and k-th columns of **X**_*p*_, **X**_*t*_, and **X**_*e*_, respectively. These columns correspond to the substructural representations of POI, PROTAC molecule, and E3 ligase. The ternary interaction can be conceptualized as first projecting representations 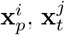, and 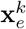 to a feature space with weight matrices **U, W**, and **V**, then learning the ternary attention map based on operation product of these projections and the weight vector **q**. This formulation of ternary interactions enables the interpretation of how substructural triplets contribute to the prediction outcomes, thereby enhancing the model’s explainability.

For the subsequent classification step, we introduce a ternary fusion layer over the attention map 𝒜 to derive the joint representation **f′**; ∈ ℝ^*κ*^. The k-th element of **f′** is computed as follows:

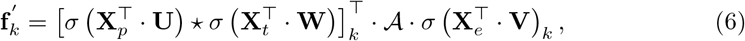

where 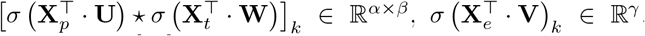, reminded that ⋆ denotes the einsum [51] *ik, jk* → *ijk*. Notably, no new learnable parameters are introduced at this layer, weight matrices **U, W**, and **V** are shared with the preceding ternary interaction layer, reducing parameter count and mitigating overfitting risk. Furthermore, we extend the single ternary interaction to a multi-head formulation by computing multiple ternary interaction maps. The final joint representation vector is obtained by summing individual heads. As the weight matrices **U, W**, and **V** are shared, each additional head introduces only one new weight **q**, ensuring parameter efficiency. In our experiments, the multi-head interaction performs better than a single one. Lastly, we apply a pooling operation on the joint representation vector to obtain a compact feature **f** = Pooling (**f′**, s), where Pooling(·) represents a one-dimensional average pooling operation with stride s, reducing the dimensionality from **f′** ∈ ℝ^*κ*^ to **f** ∈ ℝ^*κ/s*^.

In the final stage of our model, the joint representation **f** is processed through a classifier. This classifier consists of a multi-layer perceptron (MLP) classification layer, denoted as Ψ_*MLP*_, followed by a LogSoftmax activation function. The degradation prediction result **p** is computed as follows:

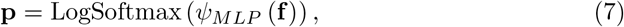

where Ψ_*MLP*_ is composed of two fully connected layers interspersed with a nonlinear activation function.

The novel ternary attention mechanism enables our model to explicitly capture and learn the intricate substructure interactions among POI, PROTAC molecule, and E3 ligase. By leveraging the learned ternary attention map, we can visualize and quantify the contribution of individual substructures to the final outcome. This approach not only aids in elucidating the underlying binding mechanisms but also significantly enhances the model’s interpretability, providing valuable insights into the complex interplay of PROTAC-mediated protein degradation.

### 4.3 Experiment settings

#### 4.3.1 Metrics

To evaluate the performance of our model and compare it with existing methods, we employed several standard metrics:

1. **Accuracy**: This metric provides the overall correct prediction rate, offering a general measure of the model’s performance.

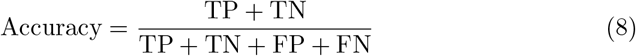

where TP, TN, FP, and FN represent True Positives, True Negatives, False Positives, and False Negatives, respectively.
2. **Area Under the Receiver Operating Characteristic curve (AUROC)**: This metric assesses the model’s ability to distinguish between classes across various threshold settings, providing a comprehensive view of classification performance.

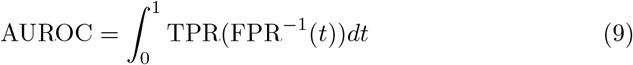

where TPR = TP / (TP + FN) is the True Positive Rate and FPR = FP / (FP + TN) is the False Positive Rate.
3. **F1 score**: The F1 score is the harmonic mean of precision and recall, offering a balanced measure of the model’s performance, particularly useful in scenarios where class imbalance may be present.

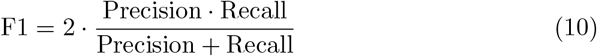

where Precision = TP / (TP + FP) and Recall = TP / (TP + FN).

These metrics collectively provide a robust evaluation for assessing the efficacy of our model in predicting degradation outcomes.

#### 4.3.2 Baselines

We compare *PROTAC-STAN* with the following three models for PROTAC degradation prediction: (1) Two classic machine learning methods, support vector matching (SVM) [32] and random forest (RF) [33] by employing the auto cross-covariance (ACC) features for the POI and E3 ligase, and the molecular access system (MACCS) keys [35] or Morgan fingerprints [34] for the PROTAC molecule following the approach described by Li et al. [17]. (2) DeepPROTACs [17] that predict PROTAC degradation using GNN to encode molecular graph of ligands and generated pockets of POI and E3 ligase, along with RNN to encode linker SMILES. The learned three features are combined with a simple concatenation to predict final degradation. For the above degradation prediction models, the SVM and RF models follow the recommended parameter settings as described in the reference paper [17], and we use the results reported in the original DeepPROTACs paper since their dataset is not publicly available.

#### 4.3.3 Implementation

*PROTAC-STAN* is implemented in Python 3.11 and PyTorch 2.1.0, along with functions from PyTorch Geometric 2.5.1, RDKit 2023.9.2, Scikit-learn 1.2.2, Pandas 2.1.1, Numpy 1.26.4, and Biopython 1.83. The training is finished under an Nvidia GeForce RTX 3090 GPU. The batch size is set to be 4 and the Adam optimizer is used with a learning rate of 0.0005. We allow the model to run for most 100 epochs with an early stop patience set to 30. The best-performing model is selected at the epoch giving the best AUROC score. In the ablation experiments, we used the same hyperparameter configuration for all experiments to ensure comparability. Each experiment was repeated 10 times, and the results reported in the table are those from the epoch that achieved the best AUROC score. For the Raw approach, the SMILESNet encoder [17] and the Ngrams encoder [13] were implemented using their original configurations. The PROTAC feature encoder consists of one linear layer with hidden dimensions [146, 64], two GCN layers with hidden dimensions [128, 128], and an edge dimension of 3. The POI and E3 ligase feature encoders consist of a PLM embedder with an embedding dimension of 1280 and two linear layers with hidden dimensions [128, 64]. The maximum allowed number of SMILES characters for PROTAC is set to 128, and the maximum permitted protein sequence length for POIs and E3 ligases is 1022. In the ternary attention module, we only employ two attention heads to provide better interpretability. The number of hidden neurons in the MLP classifier is set to 64. Part of the illustrations in this article were created using ML Visuals material. More details of the configuration are provided in Appendix B.1.

## Declarations

## Competing Interests Statement

The authors declare no competing interests.

## Data availability

All data used in this paper are publicly available and can be accessed at http://cadd.zju.edu.cn/protacdb/ for PROTAC-DB, https://github.com/PROTACs/PROTAC-STAN for PROTAC-fine dataset.

## Code availability

All code of *PROTAC-STAN* is available at https://github.com/PROTACs/PROTAC-STAN.

## Appendix A Methodologies

### A.1 Major notations

The major notations including the professional abbreviations and mathematical symbols used in this paper are provided in Table A1.

**Table A1.**
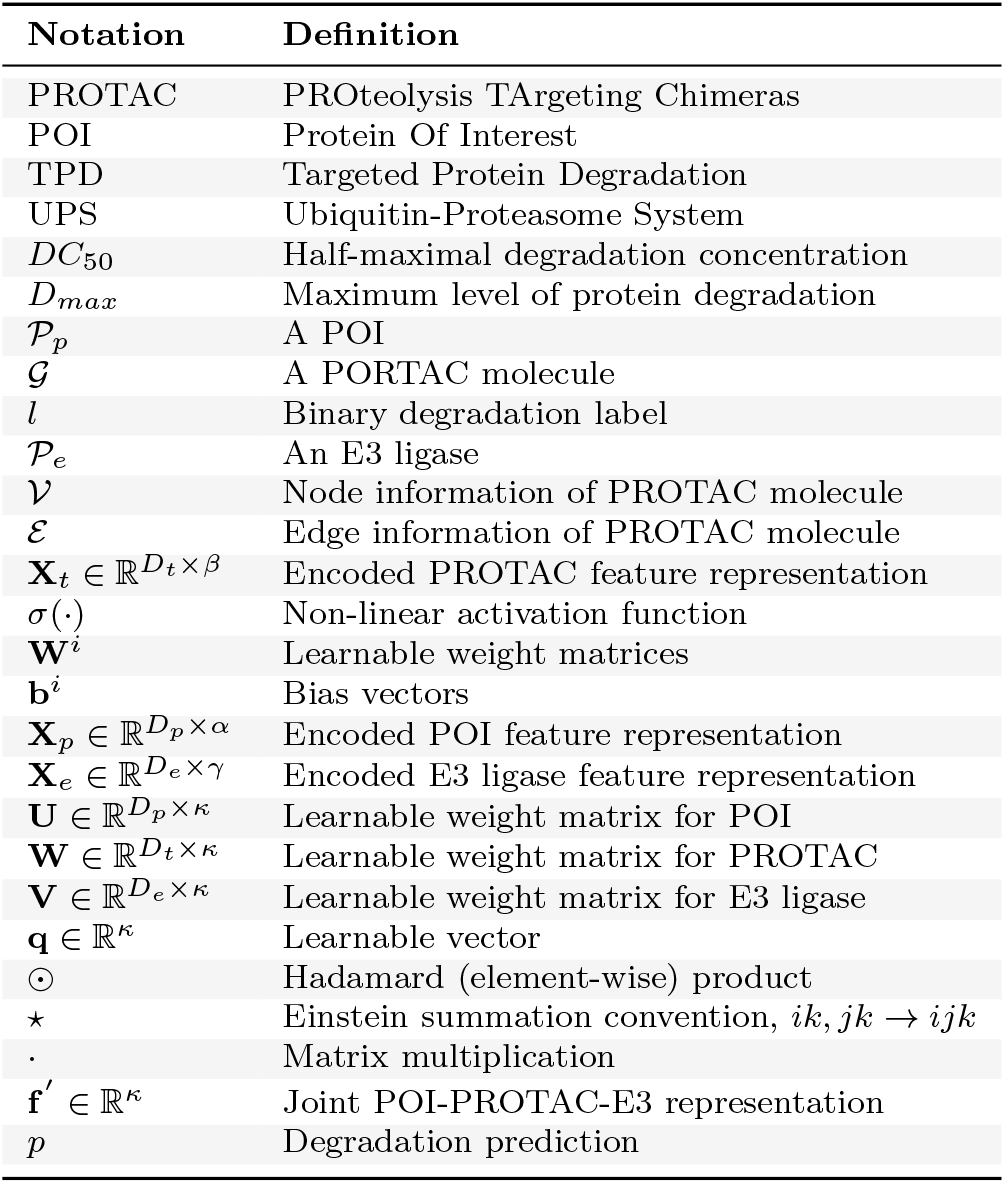
Major Notations

### A.2 Generality of TAN

#### A.2.1 Review of TAN

Given the representation of three entities inputs 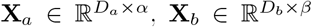, and 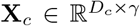, where 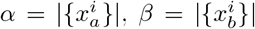, and 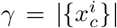 denote the number of substructures in three entities. We define TAN as a function of the inputs from three entities, parameterized by a ternary attention map, as follows:

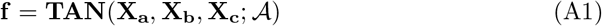

#### Ternary attention map

The ternary interaction can obtain a three-dimension ternary attention map A ∈ ℝ^*α×β×γ*^ :

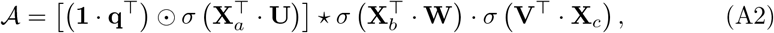

where 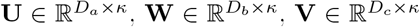 represent learnable weight matrices for three representations, respectively. **q** ∈ ℝ^*κ*^ denotes a learnable weight vector, **1** ∈ ℝ^*α*^ is a constant vector of ones. ⊙ represents the Hadamard (element-wise) product, ⋆ signifies the Einstein summation convention (einsum) [51], specially *ik, jk* → *ijk*, and · denotes standard matrix multiplication. The elements of 𝒜 quantify the interaction intensity among substructural triplets. An individual element 𝒜_*i,j,k*_ from Equation A2 can be expressed as:

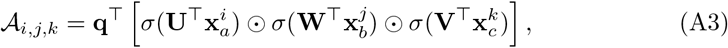

where 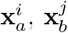, and 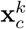 represent the i-th, j-th, and k-th columns of **X**_*a*_, **X**_*b*_, and **X**_*c*_, respectively.

#### Ternary fusion

Next, we introduce a ternary fusion layer over the attention map 𝒜 to derive the joint representation **f** ∈ ℝ^*κ*^. The k-th element of **f** is computed as follows:

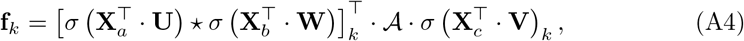

where 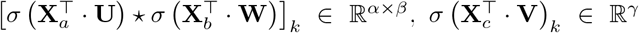. Furthermore, we extend the single ternary interaction to a multi-head formulation by computing multiple ternary interaction maps. The final joint representation vector is obtained by summing individual heads.

### A.2.2 Simplified to BAN

Bilinear Attention Network (BAN) [49] is designed for Visual Question Answering, handling two multi-channel inputs: image features **X**∈ ℝ^*N×ρ*^ and text features **Y** ∈ ℝ^*M×ϕ*^. Similar to TAN, BAN can be defined as follows:

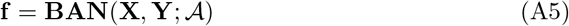

From TAN, let **X**_*a*_ ⇐ **X, X**_*b*_ ⇐ 𝕀, and **X**_*c*_ ⇐ **Y**, where 𝕀 ∈ ℝ^*D×*1^ is a vector of ones, we then obtain a two-dimension attention map A ∈ ℝ^*ρ×ϕ*^ as follows:

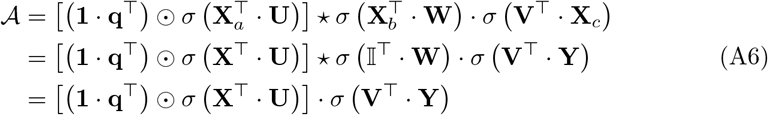

where **U** ∈ ℝ^*M×κ*^, **W** ∈ ℝ^*D×κ*^, **V** ∈ ℝ^*N×κ*^ represent learnable weight matrices for three representations, respectively. **q** ∈ ℝ^*κ*^ denotes a learnable weight vector, **1** ∈ ℝ^*ρ*^ is a constant vector of ones. An individual element 𝒜_*i,j*_ from Equation A6 can be expressed as:

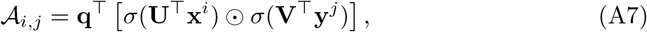

where **x**^*i*^, **y**^*j*^ represent the i-th, j-th columns of **X, Y**.

Similar to the ternary fusion, we can derive the joint representation of BAN, denoted as **f** ∈ ℝ^*κ*^, using the attention map obtained in Equation A6. The k-th element of **f** is computed as follows:

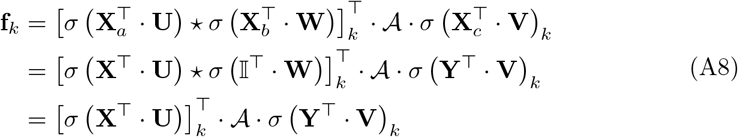

where [σ (**X**^⊤^ · **U)]** _*k*_ ∈ ℝ^*ρ*^, σ (**Y**^⊤^ · **V)**_***k***_ ∈ ℝ^*ϕ*^.

Generally, TAN can be applied to any scenario involving the interplay of three entities, providing nuanced insights into multi-entity interactions. This leads to more accurate predictions and enhanced interpretability of results. In the case of TAN and BAN, our proposed TAN offers a unified framework for attention networks with multiple inputs. Looking ahead, we may extend the Ternary Attention Network to an N-ary Attention Network, such as a quaternary attention network, in specific scenarios.

## Appendix B Implementation Details

### B.1 Hyperparemeters and configurations

Here, we demonstrate the hyperparameters and training configuration used in our methods as in Table B2.

### B.2 TAN module ablation table

The ablation results of TAN module are demonstrated in Table B3.

### B.3 Implicit *DC*_50_ and *D*_*max*_ from experimental descriptions

The explicit DC_50_ and D_*max*_ values in PROTAC-DB are relatively scarce. Upon thorough analysis of the PROTAC-DB data, we discovered that it contains partial descriptions of degradation experiments, as shown in Table B4. These primarily include

*Percent degradation* and *Assay (Percent Degradation)*. By examining the Percent degradation at the corresponding Assay DC_50_ dose, it is possible to assess the degradation efficacy. Consequently, we can derive additional degradation activity information from these experimental descriptions.

**Table B2.** Summary of hyperparameters and configurations

| Parameter | Value | Description |
| --- | --- | --- |
| *Model* |  |  |
| Max SMILES length | 128 | Maximum number of PROTAC SMILES characters |
| PROTAC feature | 146 | Dimension of input PROTAC features |
| PROTAC hidden state | 128 | Dimension of PROTAC hidden states |
| PROTAC embedding | 128 | Dimension of PROTAC embeddings |
| PROTAC edge | 3 | Dimension of PROTAC graph edges |
| Max protein length | 1022 | Maximum number of amino acids of input POIs/E3 ligases |
| Sequence embedding | 1280 | Dimension of POI/E3 Sequence embedding |
| POI/E3 hidden state | 128 | Dimension of POI/E3 hidden states |
| POI/E3 output | 64 | Dimension of POI/E3 outputs |
| Fusion feature | 192 | Dimension of fusion features |
| Multiheads | 2 | Number of attention heads |
| Dropout | [0.2, 0.2, 0.2] | Dropout ratio of each input features |
| Joint feature embedding | 192 | Dimension of joint POI-PROTAC-E3 feature embedding |
| Hidden state | 64 | Dimension of joint feature hidden states |
| Class | 2 | Number of classification classes |
| *Training* |  |  |
| Learning rate | 0.0005 | Learning rate for the optimizer |
| Batch size | 4 | Number of samples per step |
| Epochs | 100 | Number of times to iterate over the entire dataset |
| Traning ratio | 0.8 | Train dataset ratio of the entire dataset |
| Optimizer | Adam | Optimization method for updating weights |
| Loss function | Cross-entropy | Loss function for training the model |

**Table B3.**
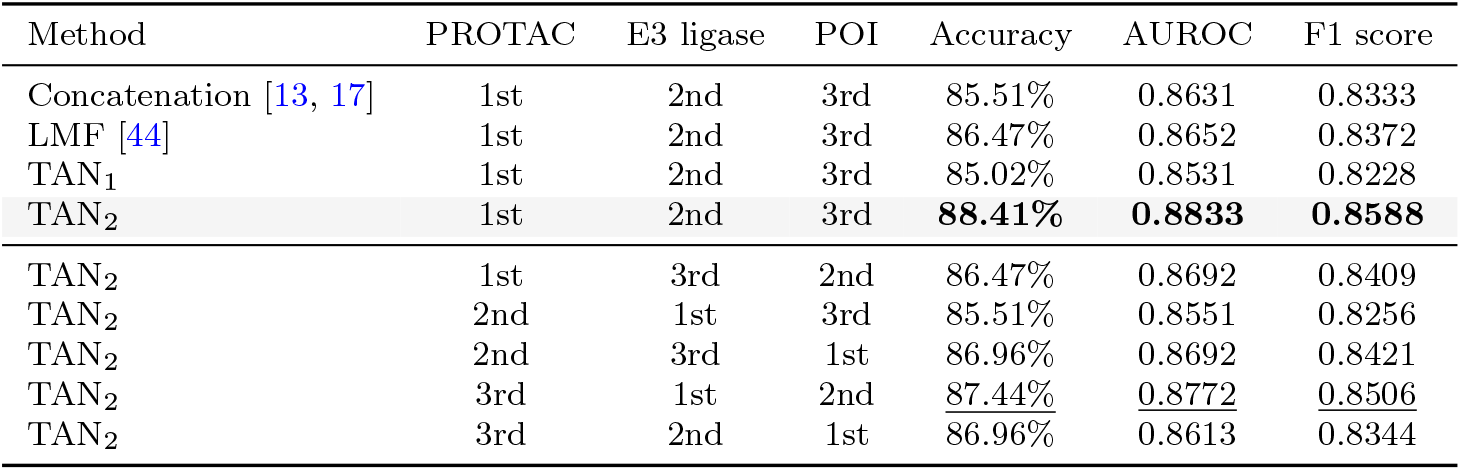
Ablation Study on TAN module. This table compares the performance of concatenation, LMF, and TAN fusion methods, investigates the effects of multi-heads of TAN, TAN_1_ for single head and TAN_2_ for two heads, and explores the impact of varying orders of input features. **Best**, Second Best.

| Method | PROTAC | E3 ligase | POI | Accuracy | AUROC | F1 score |
| --- | --- | --- | --- | --- | --- | --- |
| Concatenation [13, 17] | 1st | 2nd | 3rd | 85.51% | 0.8631 | 0.8333 |
| LMF [44] | 1st | 2nd | 3rd | 86.47% | 0.8652 | 0.8372 |
| TAN<sub>1</sub> | 1st | 2nd | 3rd | 85.02% | 0.8531 | 0.8228 |
| TAN<sub>2</sub> | 1st | 2nd | 3rd | **88.41%** | **0.8833** | **0.8588** |
| TAN<sub>2</sub> | 1st | 3rd | 2nd | 86.47% | 0.8692 | 0.8409 |
| TAN<sub>2</sub> | 2nd | 1st | 3rd | 85.51% | 0.8551 | 0.8256 |
| TAN<sub>2</sub> | 2nd | 3rd | 1st | 86.96% | 0.8692 | 0.8421 |
| TAN<sub>2</sub> | 3rd | 1st | 2nd | <u>87.44%</u> | <u>0.8772</u> | <u>0.8506</u> |
| TAN<sub>2</sub> | 3rd | 2nd | 1st | 86.96% | 0.8613 | 0.8344 |

**Table B4.**
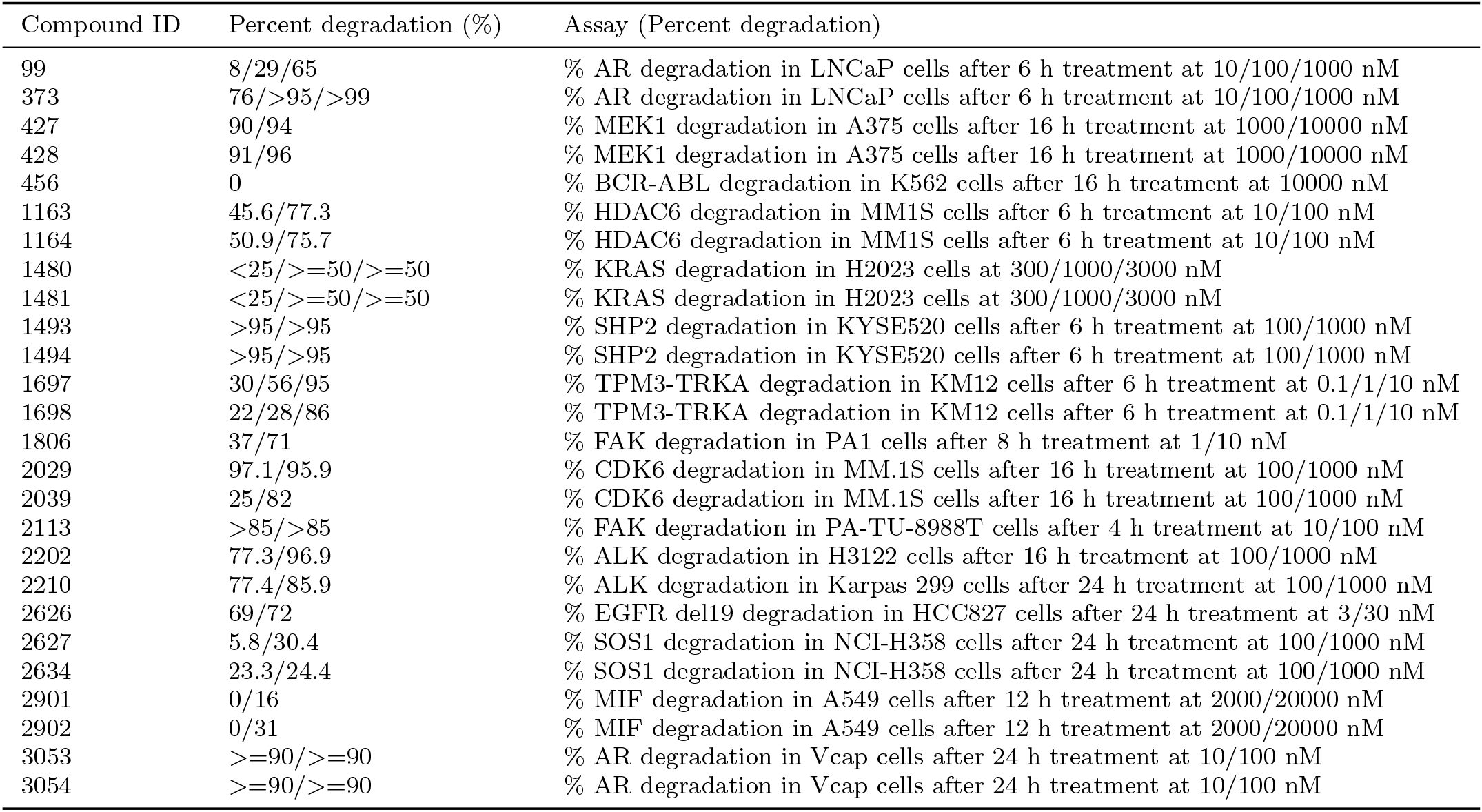
Part of degradation experimental descriptions

### B.4 Length statistics of PROTAC SMILES

We analyzed the length distribution of all PROTAC molecules based on the PROTAC table from PROTAC-DB. The results are shown in Figure B1. The average length is 136, the median length is 127, the maximum length is 285, and the minimum length is 52, so we take 128 as the maximum allowed length of PROTAC SMILES, which is the tradeoff of average length and median length.

### B.5 PROTAC properties standardization

We extracted nine physicochemical properties of PROTAC molecules from PROTAC-DB: Molecular Weight, Exact Mass, XLogP3, Heavy Atom Count, Ring Count, Hydrogen Bond Acceptor Count, Hydrogen Bond Donor Count, Rotatable Bond Count, and Topological Polar Surface Area. Figure B2 shows their raw value distributions. Due to large value disparities, we standardized each property using its mean and variance as shown in Figure B3, preserving relative characteristics while minimizing interference in experimental results.

**Fig B1.**
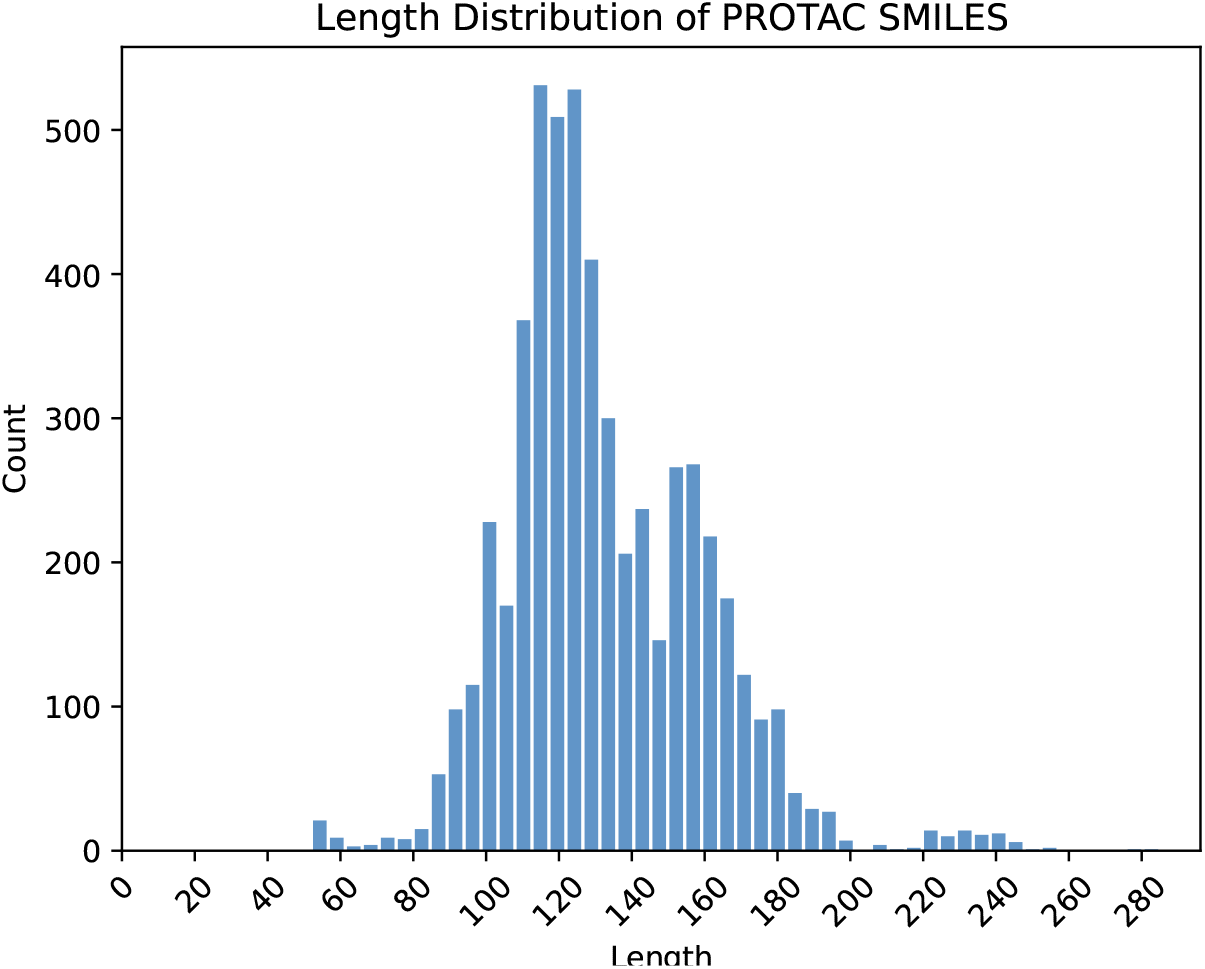
Length distribution of PROTAC SMILES

**Fig B2.**
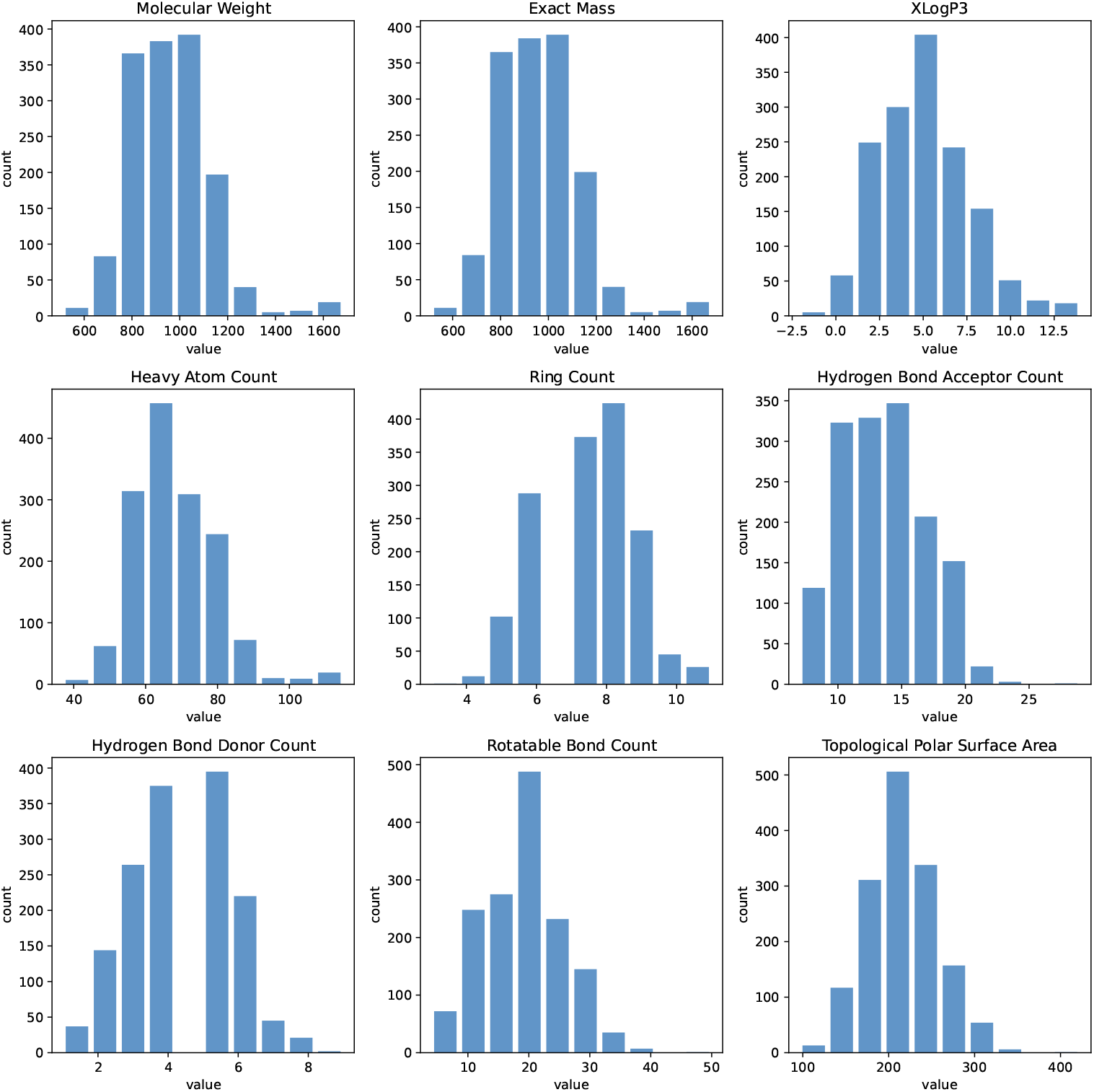
Raw PROTAC properties distribution

**Fig B3.**
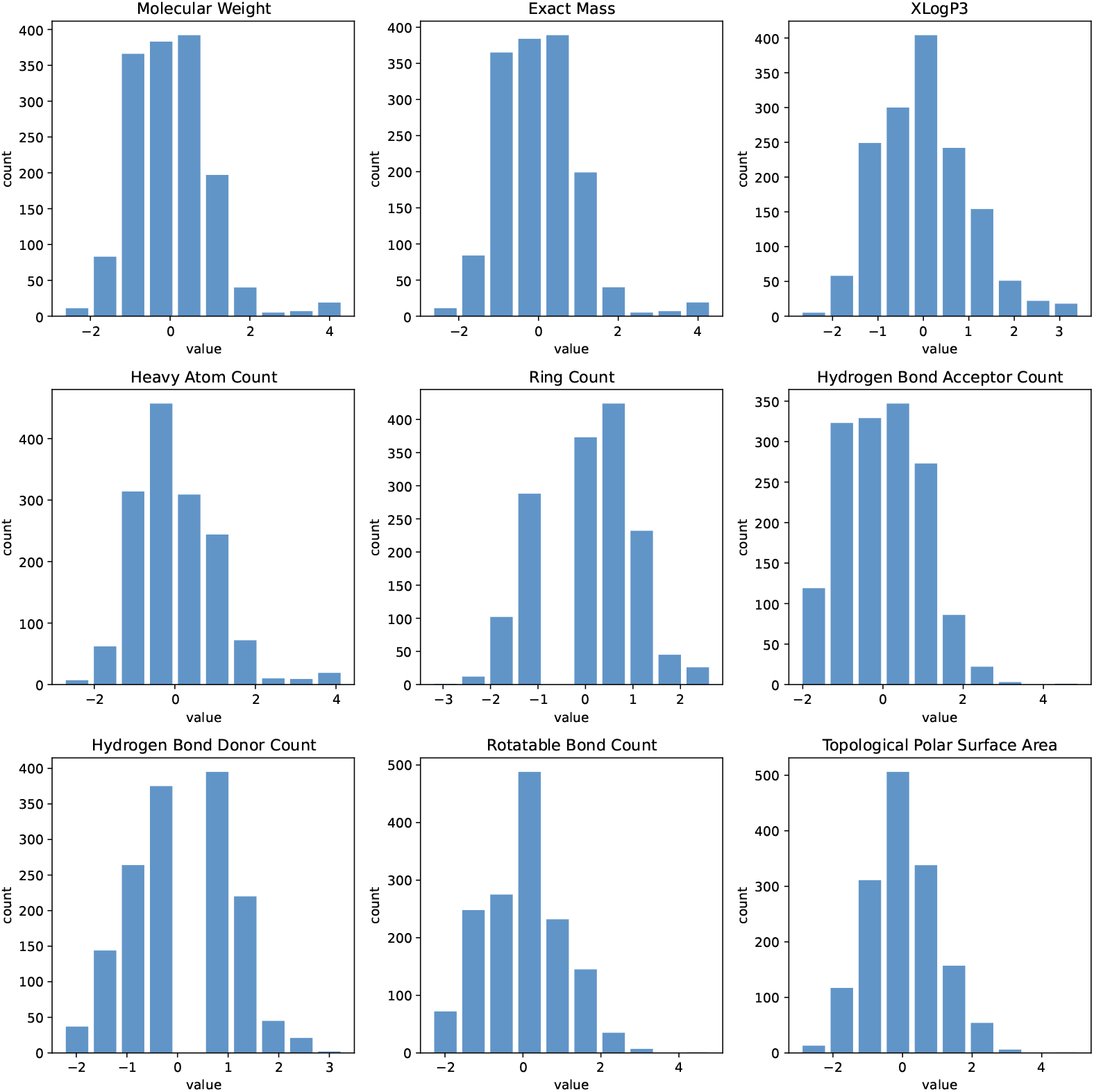
Standardized PROTAC properties distribution

**Fig B4.**
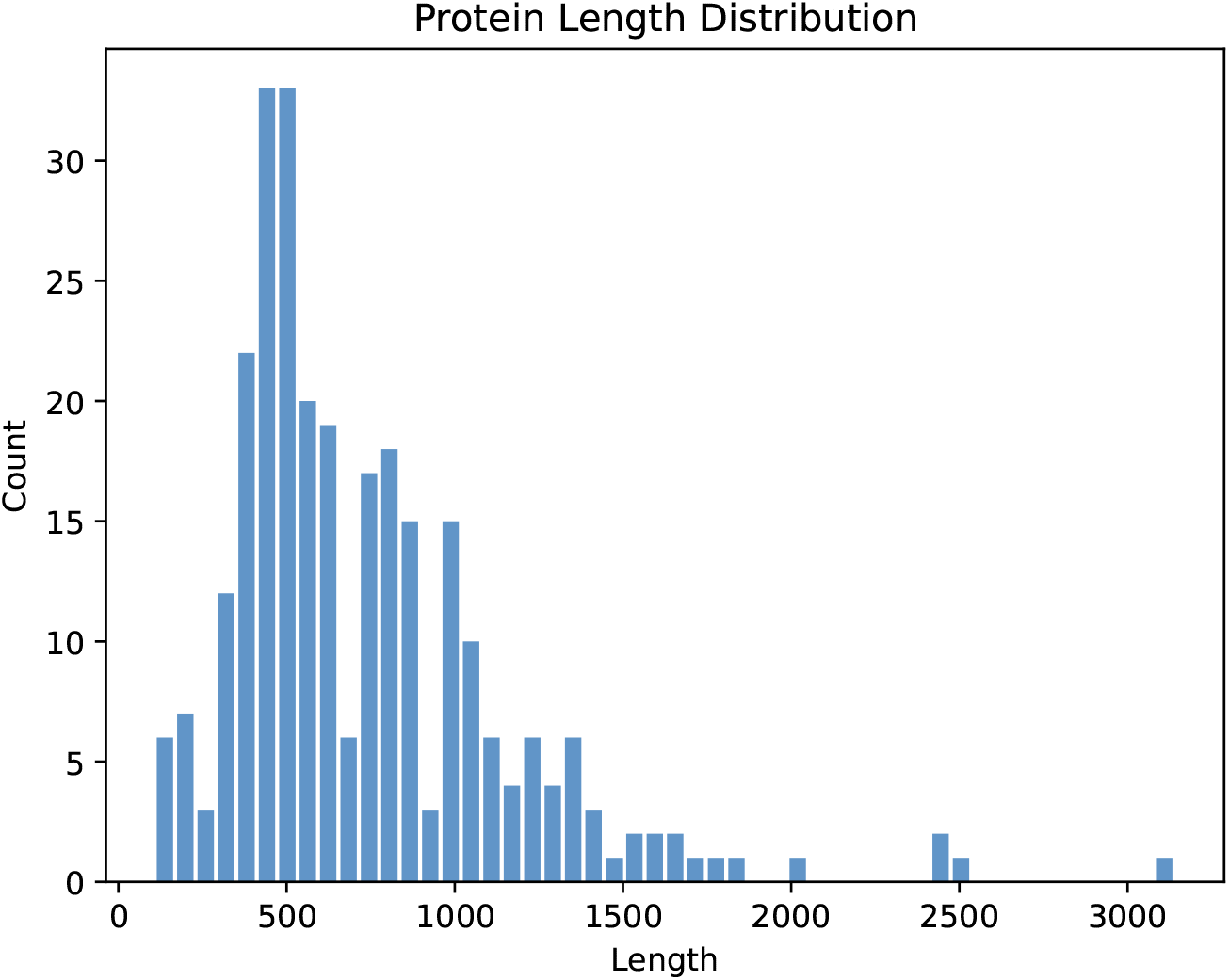
Length distribution of POI/E3 ligase

### B.5 Length statistics of POI/E3 ligase sequence

We also analyzed the length distribution of all proteins including POI and E3 ligase based on the Warhead table and E3 ligand table from PROTAC-DB. The results are as shown in Figure B4. The average length is 730, while the suggested length of the protein language model, i.e. ESM is 1022, which covers almost 80% of the proteins, so we adopt it as the maximum allowed sequence length.

## Appendix C More visualization results

### C.1 3D attention map

Here, we present additional exemplars of 3D attention map visualization, leveraging three illustrative samples from PROTAC-DB 2.0 (Compound IDs: 339, 340, and 2286). Notably, as depicted in Figure C5, a consistent trend emerges as the above observation. Specifically, the leftmost two columns exhibit pronounced patterns, whereas the figures in the right final column demonstrate weaker patterns.

### c.2 2D attention map

We extend our analysis by presenting further 2D pairwise attention map visualizations as depicted in Figure C6, derived from the above 3D attention weights. Consistent with our previous observations, the visualizations reveal a notable distinction between the pairwise attention profiles. Specifically, the pairwise attention between PROTAC and E3 ligase, as well as between PROTAC and POI, exhibits relatively high values (deep blue-purple), whereas the pairwise attention between E3 ligase and POI display relatively lower values (medium yellow).

**Fig C5.**
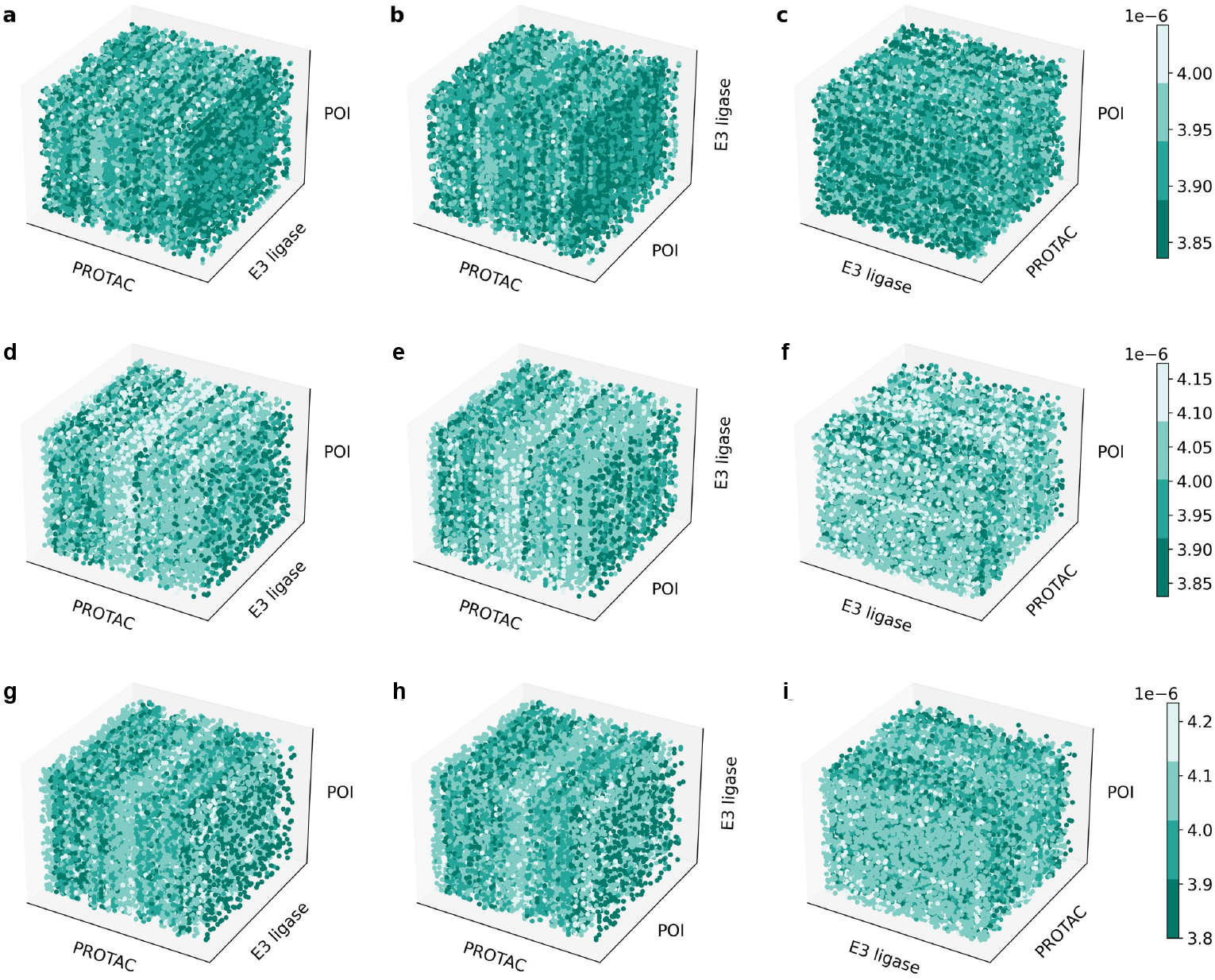
More examples of 3D attention map visualization. Sample 339 (a-c), sample 340 (d-f), and sample 2286 (g-i). Three figures in each row are front view, top view, and side view, respectively.

**Fig C6.**
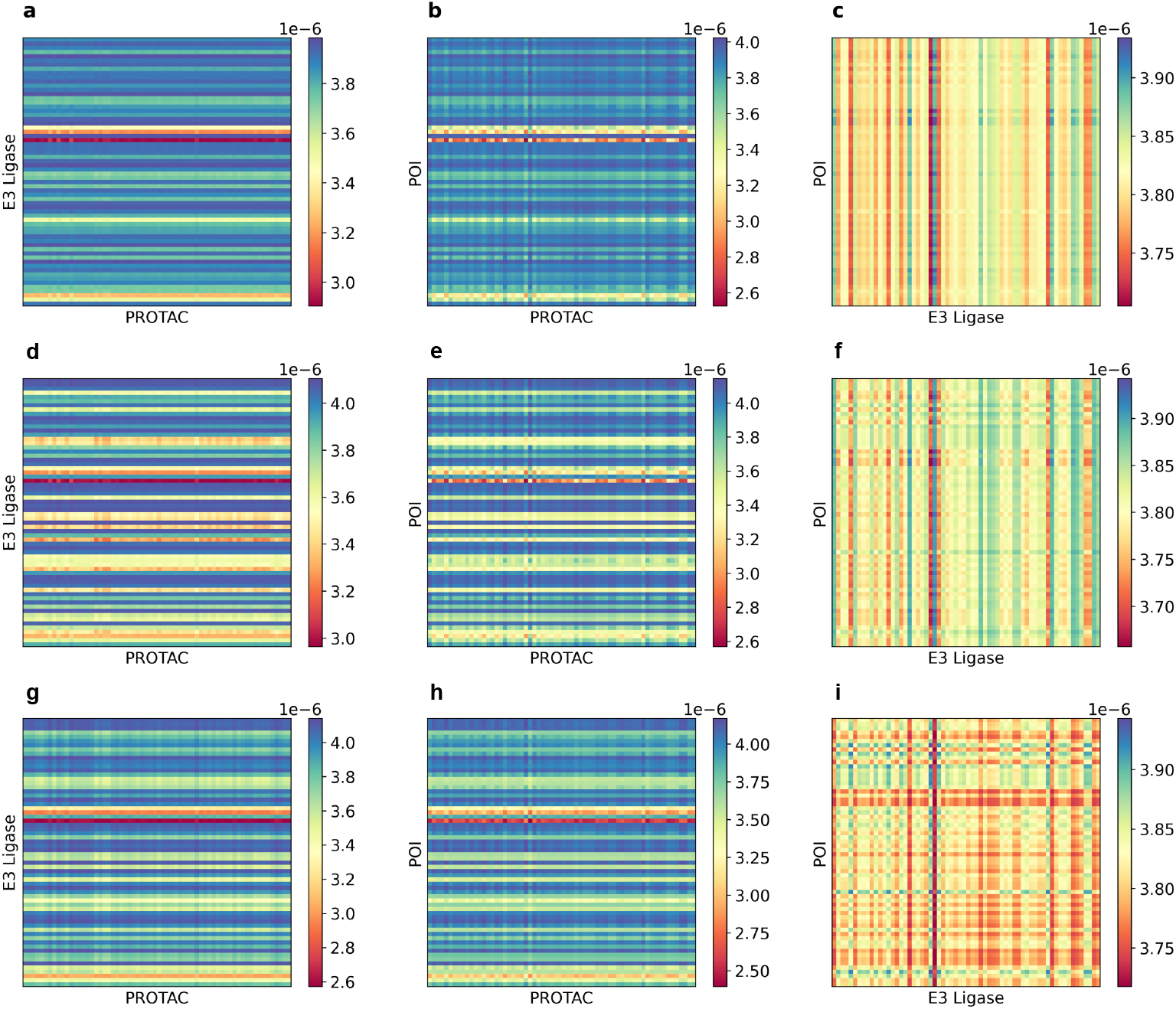
More examples of 2D attention map visualization. Sample 339 (a-c), sample 340 (d-f), and sample 2286 (g-i). Three figures in each row are pairwise attention maps of PROTAC-E3 ligase, PROTAC-POI, and E3 ligase-POI, respectively.

### c.3 2D PROTAC molecule

We continue to map attention weights onto PROTAC molecules and 3D complexes. The visualized 2D PROTAC molecules with weighted atoms are demonstrated in Figure C7.

**Fig C7.**
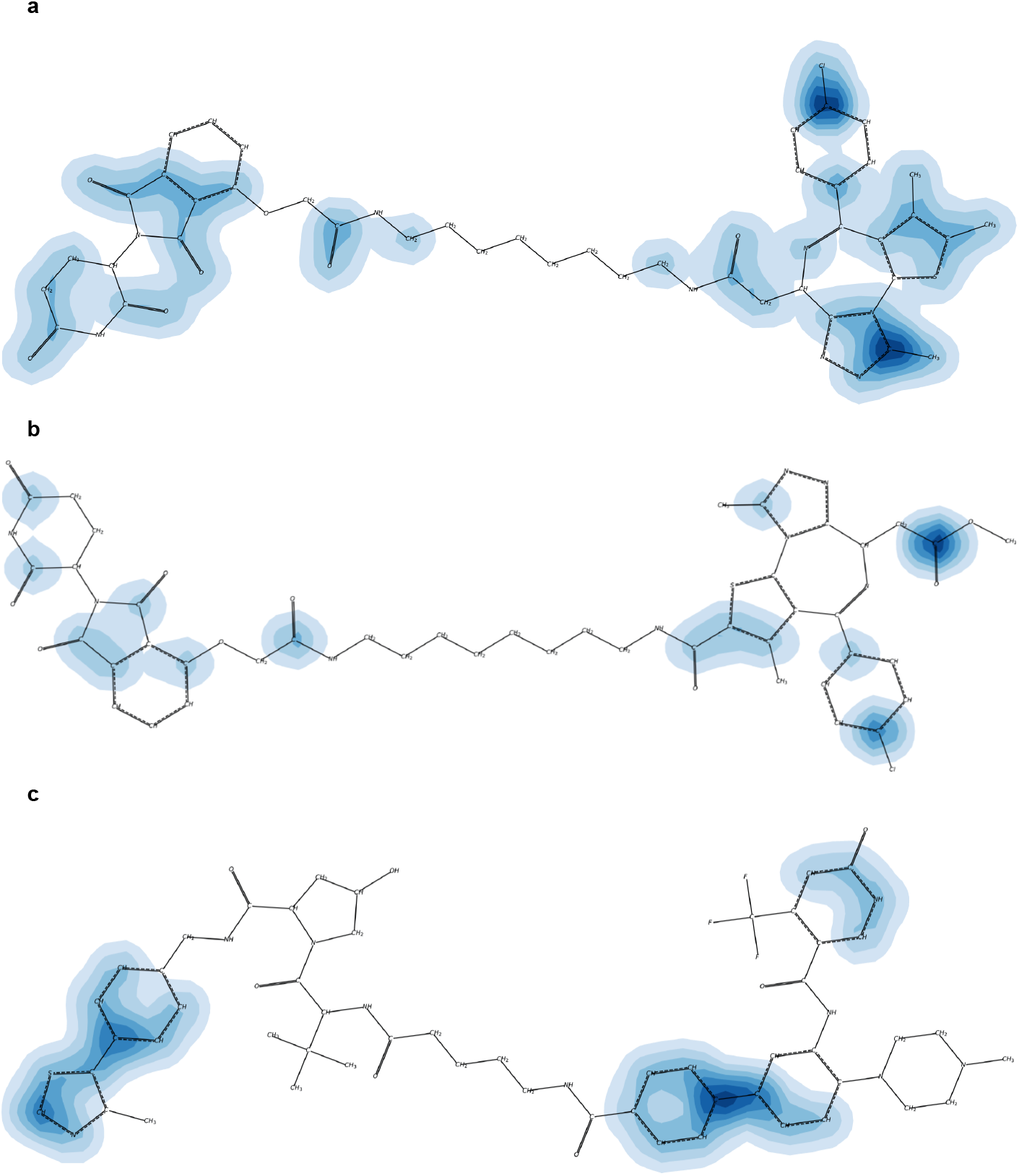
More examples of weighted 2D PROTAC molecule. a. Compound 339. **b**. Compound 340. **c**. Compound 2286.

**Fig C8.**
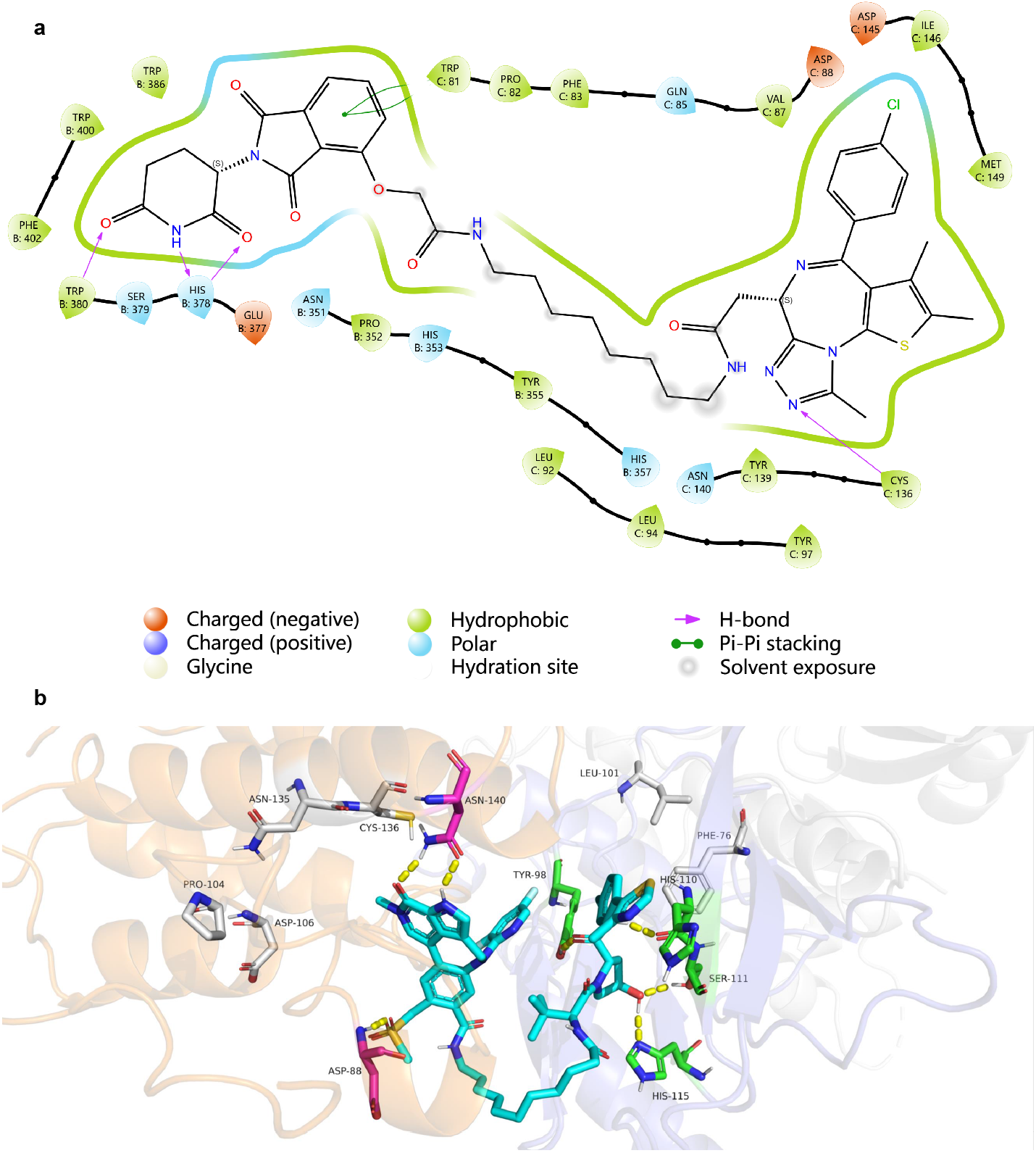
2D interaction and 3D Complex visualization for example PDB 7Q2J. **a**, 2D interactions of PROTAC and protein. There are mainly hydrogen bonding and *π*-*π* stacking interactions between PROTACs and proteins. **b**.3D pocket visualization. The top important weights in gray, and marked the actual interacting residues in green, overlapped residues in magenta, hydrogen bonds in yellow, and *π*-*π* stacking interactions in green. The blue-purple part in the background represents E3 ligase, while the orange part represents POI.

### c.4 2D interaction and 3D Complex

The samples with 3D crystal structures are scarce. Consequently, we visualize one more example,

## Notes

### Competing Interest Statement

The authors have declared no competing interest.

## References

[1] Zhao, L., Zhao, J., Zhong, K., Tong, A. & Jia, D. Targeted protein degradation: Mechanisms, strategies and application. Signal Transduction and Targeted Therapy 7, 113 (2022).

[2] Békés, M., Langley, D. R. & Crews, C. M. PROTAC targeted protein degraders: The past is prologue. Nature Reviews Drug Discovery 21, 181–200 (2022).

[3] Chirnomas, D., Hornberger, K. R. & Crews, C. M. Protein degraders enter the clinic — a new approach to cancer therapy. Nature Reviews Clinical Oncology 20, 265–278 (2023).

[4] Lai, A. C. & Crews, C. M. Induced protein degradation: An emerging drug discovery paradigm. Nature Reviews Drug Discovery 16, 101–114 (2017).

[5] Burslem, G. M. & Crews, C. M. Proteolysis-Targeting Chimeras as Therapeutics and Tools for Biological Discovery. Cell 181, 102–114 (2020).

[6] Pereira, G. P. et al. Rational Prediction of PROTAC-Compatible Protein–Protein Interfaces by Molecular Docking. Journal of Chemical Information and Modeling 63, 6823–6833 (2023).

[7] Testa, A. et al. 3-Fluoro-4-hydroxyprolines: Synthesis, Conformational Analysis, and Stereoselective Recognition by the VHL E3 Ubiquitin Ligase for Targeted Protein Degradation. Journal of the American Chemical Society 140, 9299–9313 (2018).

[8] Vamathevan, J. et al. Applications of machine learning in drug discovery and development. Nature Reviews Drug Discovery 18, 463–477 (2019).

[9] Gharbi, Y. & Mercado, R. A Comprehensive Review of Emerging Approaches in Machine Learning for De Novo PROTAC Design (2024). 2406.16681.

[10] Xie, L. & Xie, L. Elucidation of genome-wide understudied proteins targeted by PROTAC-induced degradation using interpretable machine learning. PLOS Computational Biology 19, e1010974 (2023817).

[11] Zhang, W. et al. Machine Learning Modeling of Protein-intrinsic Features Predicts Tractability of Targeted Protein Degradation. Genomics, Proteomics & Bioinformatics 20, 882–898 (2022).

[12] García Jiménez, D. et al. Designing Soluble PROTACs: Strategies and Preliminary Guidelines. Journal of Medicinal Chemistry 65, 12639–12649 (2022).

[13] Nori, D., Coley, C. W. & Mercado, R. De novo PROTAC design using graph-based deep generative models (2022). 2211.02660.

[14] Askr, H. et al. Deep learning in drug discovery: An integrative review and future challenges. Artificial Intelligence Review 56, 5975–6037 (2023).

[15] Catacutan, D. B., Alexander, J., Arnold, A. & Stokes, J. M. Machine learning in preclinical drug discovery. Nature Chemical Biology 1–14 (2024).

[16] Guo, J. et al. Link-INVENT: Generative linker design with reinforcement learning. Digital Discovery 2, 392–408 (2023).

[17] Li, F. et al. DeepPROTACs is a deep learning-based targeted degradation predictor for PROTACs. Nature Communications 13, 7133 (2022).

[18] Li, B., Ran, T. & Chen, H. 3D based generative PROTAC linker design with reinforcement learning. Briefings in Bioinformatics bbad323 (2023).

[19] Kao, C.-T., Lin, C.-T., Chou, C.-L. & Lin, C.-C. Fragment Linker Prediction Using the Deep Encoder-Decoder Network for PROTACs Drug Design. Journal of Chemical Information and Modeling 63, 2918–2927 (2023).

[20] Igashov, I. et al. Equivariant 3D-conditional diffusion model for molecular linker design. Nature Machine Intelligence 6, 417–427 (2024).

[21] Mslati, H., Gentile, F., Pandey, M., Ban, F. & Cherkasov, A. PROTACable Is an Integrative Computational Pipeline of 3-D Modeling and Deep Learning To Automate the De Novo Design of PROTACs. Journal of Chemical Information and Modeling (2024).

[22] Zheng, S. et al. Accelerated rational PROTAC design via deep learning and molecular simulations. Nature Machine Intelligence 4, 739–748 (2022).

[23] Yang, Y. et al. SyntaLinker: Automatic fragment linking with deep conditional transformer neural networks. Chemical Science 11, 8312–8322 (2020).

[24] Tan, Y. et al. DRlinker: Deep Reinforcement Learning for Optimization in Fragment Linking Design. Journal of Chemical Information and Modeling 62, 5907–5917 (2022).

[25] Schenone, M., Dančík, V., Wagner, B. K. & Clemons, P. A. Target identification and mechanism of action in chemical biology and drug discovery. Nature Chemical Biology 9, 232–240 (2013).

[26] Ermondi, G., Garcia-Jimenez, D. & Caron, G. PROTACs and Building Blocks: The 2D Chemical Space in Very Early Drug Discovery. Molecules 26, 672 (2021).

[27] Jiménez-Luna, J., Grisoni, F. & Schneider, G. Drug discovery with explainable artificial intelligence. Nature Machine Intelligence 2, 573–584 (2020).

[28] Kipf, T. N. & Welling, M. Semi-Supervised Classification with Graph Convolutional Networks (2017). 1609.02907.

[29] Lin, Z. et al. Evolutionary-scale prediction of atomic-level protein structure with a language model. Science (2023).

[30] Zhang, Z. et al. Structure-Informed Protein Language Model (2024). 2402.05856.

[31] Weng, G. et al. PROTAC-DB 2.0: An updated database of PROTACs. Nucleic Acids Research 51, D1367–D1372 (2023).

[32] Cortes, C. & Vapnik, V. Support-vector networks. Machine Learning 20, 273–297 (1995).

[33] Ho, T. K. Random decision forests, Vol. 1, 278–282 vol.1 (1995).

[34] Morgan, H. L. The generation of a unique machine description for chemical structures-a technique developed at chemical abstracts service. Journal of Chemical Documentation 5, 107–113 (1965).

[35] Durant, J. L., Leland, B. A., Henry, D. R. & Nourse, J. G. Reoptimization of MDL keys for use in drug discovery. Journal of Chemical Information and Computer Sciences 42, 1273–1280 (2002).

[36] Weiss, D. R. et al. On ternary complex stability in protein degradation: In silico molecular glue binding affinity calculations. Journal of Chemical Information and Modeling 63, 2382–2392 (2023).

[37] Roy, M. J. et al. SPR-measured dissociation kinetics of PROTAC ternary complexes influence target degradation rate. ACS Chemical Biology 14, 361–368 (2019).

[38] Hughes, S. J. & Ciulli, A. Molecular recognition of ternary complexes: A new dimension in the structure-guided design of chemical degraders. Essays in Biochemistry 61, 505–516 (2017).

[39] Ge, J. et al. PROTAC-DB 3.0: An updated database of PROTACs with extended pharmacokinetic parameters. Nucleic Acids Research gkae768 (2024).

[40] Dragovich, P. S. et al. Antibody-mediated delivery of chimeric BRD4 degraders. Part 2: Improvement of in vitro antiproliferation activity and in vivo antitumor efficacy. Journal of Medicinal Chemistry 64, 2576–2607 (2021).

[41] Yu, X. et al. A selective WDR5 degrader inhibits acute myeloid leukemia in patient-derived mouse models. Science Translational Medicine 13, eabj1578 (2021).

[42] Berman, H. M. et al. The protein data bank. Nucleic Acids Research 28, 235–242 (2000).

[43] Dong, Y. et al. Characteristic roadmap of linker governs the rational design of PROTACs. Acta Pharmaceutica Sinica B 14, 4266–4295 (2024).

[44] Liu, Z. et al. Gurevych, I. & Miyao, Y. (eds) Efficient Low-rank Multimodal Fusion With Modality-Specific Factors. (eds Gurevych, I. & Miyao, Y.) Proceedings of the 56th Annual Meeting of the Association for Computational Linguistics (Volume 1: Long Papers), 2247–2256 (Melbourne, Australia, 2018).

[45] Riching, K. M. et al. Quantitative Live-Cell Kinetic Degradation and Mechanistic Profiling of PROTAC Mode of Action. ACS Chemical Biology 13, 2758–2770 (2018).

[46] Sterling, T. & Irwin, J. J. ZINC 15 – Ligand Discovery for Everyone. Journal of Chemical Information and Modeling 55, 2324–2337 (2015).

[47] The UniProt Consortium. UniProt: The Universal Protein Knowledgebase in 2023. Nucleic Acids Research 51, D523–D531 (2023).

[48] Kim, J.-H. et al. Hadamard Product for Low-rank Bilinear Pooling (2017). 1610.04325.

[49] Kim, J.-H., Jun, J. & Zhang, B.-T. Bilinear Attention Networks (2018). 1805.07932.

[50] Bai, P., Miljković, F., John, B. & Lu, H. Interpretable bilinear attention network with domain adaptation improves drug–target prediction. Nature Machine Intelligence 5, 126–136 (2023).

[51] Einstein, A. Die Grundlage der allgemeinen Relativitätstheorie. Annalen der Physik 354, 769–822 (1916).

